# *In Vivo* Blunt-End Cloning Through CRISPR/Cas9-Facilitated Non-Homologous End-Joining

**DOI:** 10.1101/019570

**Authors:** Jonathan M. Geisinger, Sören Turan, Sophia Hernandez, Laura P. Spector, Michele P. Calos

## Abstract

The ability to precisely modify the genome in a site-specific manner is extremely useful. The CRISPR/Cas9 system facilitates precise modifications by generating RNA-guided double-strand breaks. We demonstrate that guide RNA pairs generate deletions that are repaired with a high level of precision by non-homologous end-joining in mammalian cells. We present a method called knock-in blunt ligation for exploiting this excision and repair to insert exogenous sequences in a homology-independent manner without loss of additional nucleotides. We successfully utilize this method in a human immortalized cell line and induced pluripotent stem cells to insert fluorescent protein cassettes into various loci, with efficiencies up to 35.8% in HEK293 cells. We also present a version of Cas9 fused to the FKBP12-L106P destabilization domain for investigating repair dynamics of Cas9-induced double-strand breaks. Our *in vivo* blunt-end cloning method and destabilization-domain-fused Cas9 variant increase the repertoire of precision genome engineering approaches.

## Introduction

The ability to make precise double-strand breaks (DSBs) in the genome is extremely useful for genome engineering, as this ability can facilitate directed changes in the genome for applications ranging from studying a gene to engineering an entire biosynthetic pathway. The most popular tools for making DBSs are currently zinc finger nucleases (ZFNs), transcription activator-like effector nucleases (TALENs), and the clustered regularly interspaced short palindromic repeat (CRISPR)/Cas9 system. Of these three, the CRISPR/Cas9 system offers the greatest ease of use.

The CRISPR/Cas9 system, a component of the bacterial RNA-mediated adaptive immune system, consists of transcribed guide-RNAs derived from integrated fragments of phage and plasmid DNA that direct the Cas9 RNA-guided DNA endonuclease to targets containing complementary sequences to the CRISPR (Barrangou *et al*., 2007; Jinek *et al*., 2012). The complex requires a protospacer adjacent motif (PAM) downstream of the target sequence to begin binding (Sternberg, Redding *et al*., 2014). Upon binding, Cas9 generates a blunt DSB three to four bases upstream of the PAM through the use of RuvC-like and HNH nuclease domains (Jinek *et al*., 2012).

Recently, the CRISPR/Cas9 system from *Streptococcus pyogenes* has been adapted for genome editing (Jinek *et al*., 2012; Mali, Yang *et al*., 2013; Jinek *et al.,* 2013; Cong *et al*., 2013), and has been successfully used in many eukaryotes (Friedland *et al*., 2013; Kim, Ishidate *et al*., 2014; Zhang, Koolhass, and Schnorrer, 2014; Gratz, Ukken *et al*., 2014; DiCarlo *et al*., 2013; Harel *et al*., 2015), as well as in disease and therapeutic modeling (Xue, Chen, Yin *et al*., 2014; Heckl *et al*., 2014; Platt, Chen *et al*., 2014; Ousterout *et al*., 2015; Hu, Kaminski *et al*., 2014; Wang and Quake, 2014). However, the spectre of off-target cleavage may present significant challenges to the use of CRISPR/Cas9 in these applications. The chimeric single-guide RNA (sgRNA) not only tolerates mismatches, insertions, and deletions throughout its target sequence, particularly outside of the seed sequence, but may also display cell type-specific off-target effects (Fu *et al*., 2013; Hsu *et al*., 2013; Pattanayak *et al*., 2013; Cho *et al*., 2014; Lin *et al*., 2014; Fu *et al*., 2014; Wu *et al*., 2014; Kuscu, Arslan *et al*., 2014; Cencic, Miura *et al*., 2014). Contrary to these observations, off-target cleavage in murine embryos and human induced pluripotent and embryonic stem cells (hiPSCs and ESCs) has been reported to be extremely rare (Yang, Wang *et al*., 2013; Veres *et* al., 2013; Xie *et al*., 2014; Yang, Grishin, Wang *et al*., 2014).

This discrepancy may be explained by the underlying repair mechanism of CRISPR/Cas9-induced DSBs. Typically, these DSBs are thought to be repaired by error-prone NHEJ, similarly to those generated by the *Fok*I domains of TALENs and ZFNs (Jinek *et al*., 2013; Mali, Yang *et al*., 2013; Cong *et al*., 2013). However, Cas9 makes blunt DSBs, whereas FokI dimers make overhangs. Thus, these blunt, chemically unmodified DSBs could be repaired by the precise NHEJ pathway, which has been previously demonstrated using the Tn5 transposon system, which also makes blunt DSBs (van Heemst *et al*., 2004). Indeed, precise NHEJ has been observed to resolve the majority of CRISPR/Cas9-induced translocation events in human immortalized cells (Ghezraoui *et al*., 2014). Additionally, the predominant use of precise NHEJ may explain the relatively low level of error-prone editing observed in hiPSCs (Mali, Yang *et al*., 2013).

There appears to be a large amount of variation in sgRNA efficiency, as measured by error-prone repair between different cell types (Mali, Yang *et al*., 2013). Because Cas9-mediated cleavage generates DSBs with blunt ends, we hypothesized that the error-prone NHEJ is a by-product of the propensity of Cas9 to remain bound to its target DNA sequence after cleavage (Sternberg, Redding *et al*., 2014). Thus, precise NHEJ should be the contributing factor to resolving a significant percentage of Cas9-mediated DSBs, potentially explaining the large degree in variation. We further hypothesized that precise NHEJ could be exploited to increase the specificity of Cas9-based genome editing, which has also been suggested based on experiments in zebrafish (Auer *et al*., 2014). Additionally, we sought to restrict the window of Cas9 activity to limit both the amount of potential off-target activity and DNA-damage-associated cell cycle delay, which has been demonstrated in budding yeast (Demeter *et al*., 2000).

Here, we demonstrate that, by generating two DSBs with two different sgRNAs at the same time, both large and small precise deletions can be generated, that these DSBs can be exploited to knock in various expression cassettes and gene modifications through precise NHEJ, and that this strategy is applicable in both human and mouse cells. Additionally, we present a destabilized variant of Cas9, which we originally designed to facilitate greater control over genome modification, but which was repurposed for investigation of the repair dynamics and outcomes of Cas9-mediated DSBs.

## Results

### Patterns of Excision and Repair Resulting from Paired PAM Orientations

Because Cas9 generates blunt DSBs, we hypothesized that these blunt ends would be preferentially repaired by precise NHEJ. To test this hypothesis, we chose to generate two DSBs within the same locus rather than one because precise NHEJ mediated by one individual sgRNA would be indistinguishable from uncleaved DNA at the sequence level. However, if a cell received two sgRNA vectors expressing two different sgRNAs, there would be a high probability of excision of the intervening sequence between the paired sgRNA targets, allowing for religation of the genomic junctions and subsequent detection and analysis via PCR and Sanger sequencing (Figure 1A). Others (Cong *et al*., 2013; Mali, Yang, *et al*., 2013; Canver, Bauer *et al*., 2014) have also observed this resulting excision and religation. This excision may occur because Cas9 appears to bind the PAM side of the DSB less strongly after cleavage (Sternberg, Redding *et al*., 2014). A high level of precision has also been observed in translocations generated by two sgRNAs (Ghezraoui *et al*., 2014). However, the repair consequences of paired PAM orientations resulting from using two sgRNAs simultaneously have not been explored. For pairs of sgRNAs, there are four possible orientations: 1) PAMs Out, 2) PAMs In, 3) PAMs 5’, and 4) PAMs 3’ (Figure 1B). It is important to note that PAMs 5’ and PAMs 3’ are functionally equivalent at first glance. Because Cas9 can cleave at either the third or fourth base upstream of the PAM, precise rejoining can occur with 0, +1, or -1 bases on each of the repaired genomic ends, depending on which PAM orientation is used. As such, each orientation has four possible outcomes for precise repair (detailed in Figure 1B and shown as 5’sgRNA bases/ 3’ sgRNA bases). To this end, we generated multiple sgRNAs against the 3’UTRs of human MYOD1, PAX3, PAX7, and mouse Utrophin (Utrn) to examine the repair patterns and degree of precision utilized by paired PAM orientations (Figure 1C-E).

**Figure 1.**
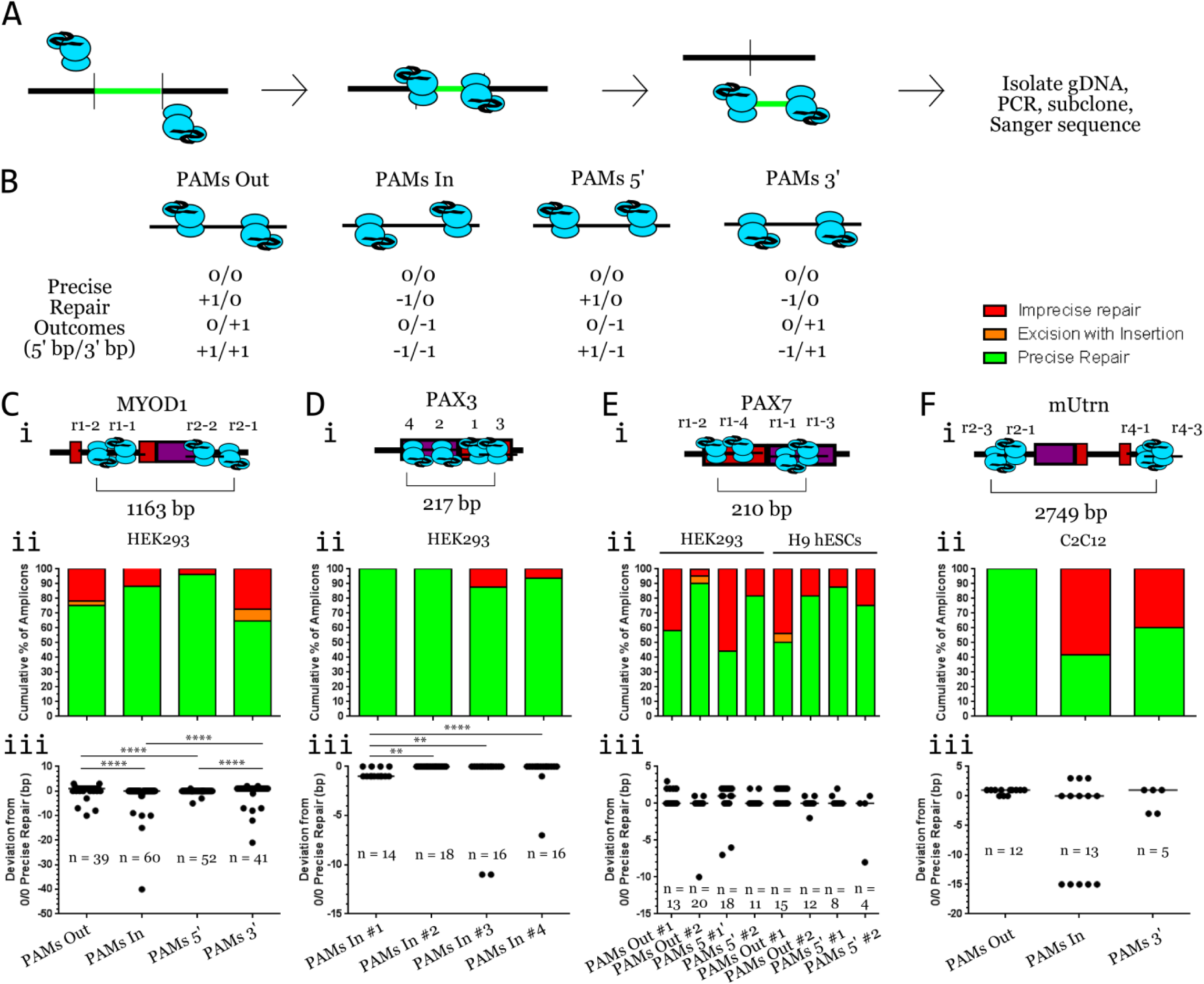
Analysis of deletion repair generated by paired sgRNAs. (A) Schematic showing excision of sequence between two sgRNAs followed by how deletion repair was analyzed. (B) Each paired PAM orientation has four possible outcomes for precise repair (depicted below each orientation) based on SpCas9’s ability to cleave at the third or the fourth base upstream of the PAM. (C-F) (i) Schematic diagrams of the targeted locus with sgRNA targets depicted as sgRNA/Cas9 complexes. Additionally, the largest deletion size for each locus is depicted. Red indicates exonic sequence and purple indicates 3’UTR. (ii) Graphs depicting the cumulative percentage of precise (green), excision+insertion (orange), and imprecise (red) repair observed in sequenced amplicons for each tested sgRNA pair. (iii) Dot plots of deletion-containing amplicons described in sub-panel ii. Each dot represents the sum of bases beyond 0/0 precise repair. The numbers of amplicons analyzed for each sgRNA pair appear on the subpanel iii graphs. Data from panel iii graphs were analyzed with the Kruskal-Wallis test followed by post-hoc Dunn’s multiple comparisons test. For MYOD1, PAX3, and mUtrn, all sgRNA pairs were compared against all pairs. For PAX7, each pair was only compared between cell types. **** = P < 0.0001; ** = P < 0.004.

To date, the largest study of repair patterns of paired PAM-mediated excisions focused only on the PAMs 3’ orientation (Canver, Bauer *et al*., 2014). In our study, we first sought to examine if there were any large differences in repair patterns between all four orientations. Each pair was tranfected separately into HEK293 cells. In examining all four paired PAM orientations at the MYOD1 3’UTR in HEK293 cells, we observed a remarkably high level of precision of NHEJ repair as indicated by the high frequency (64.71-96.15% of observed amplicons) of precise repair, regardless of orientation (Figure 1Ci and ii; for sequences, see Figure 1-figure supplement 1). PAMs 5’ showed the highest frequency of precise repair, reaching 96.15%, whereas PAMs 3’ showed the lowest frequency of precise repair (Figure 1Cii). Imprecise repair was mainly limited to small deletions and occasional inclusion of bases that should have been excised. PAMs Out and PAMs 3’ displayed a deviation from 0/0 precise repair, with both possessing a mostly 0/+1 precise repair pattern, which may be the result of them sharing the same 3’ sgRNA (Figure 1Ciii). Interestingly, the predominance of 0/+1 precise repair observed with PAMs 3’ appears to be due to 0/0 repair generating a perfect target site for the 5’ sgRNA in this pair. Comparing the deviations from 0/0 precise repair between PAM orientations revealed that there was a significant difference between distributions (*P* < 0.0001; Kruskal-Wallis test). *Post hoc* multiple comparisons revealed no significant difference between PAMs Out and PAMs 3’ or between PAMs In and PAMs 5’ (*P* > 0.9999 for both comparisons), but that all other comparisons between orientations were significantly different (*P* < 0.0001; Dunn’s multiple comparisons test for all comparisons). Additionally, we observed only one stereotypical error-prone event (Figure 1-figure supplement 1). These findings suggest that there are no inherent differences in repair precision between paired PAM orientations, the differences are dependent on individual sgRNAs, and NHEJ repair of CRISPR/Cas9-induced DSBs is not inherently error-prone.

Having observed that all four orientations were capable of facilitating precise repair at one locus, we next asked if different paired PAMs in the same orientation would display the same degree of repair precision at the same locus. Thus, we utilized four pairs of sgRNAs in the PAMs In orientation at the PAX3 3’UTR in HEK293 cells (Figure 1Di). We observed that the vast majority of sequenced deletion-containing amplicons for all four pairs were remarkably precise, with the frequency of precise repair ranging from 87.5% to 100% (Figure 1Dii; Figure 1-figure supplement 2). Interestingly, we did not observe insertions in any of these sequenced amplicons. We observed that there was a significant difference between the distributions of deviations (*P* < 0.0001; Kruskal-Wallis test), but that only PAMs In #1 displayed a noticeable deviation from 0/0 precise repair. The majority of these sequenced amplicons were of the 0/-1 precise repair variety, which was found to be significantly different than that of the other three pairs (P < 0.004 for PAMs In #1 versus each other pair; *post hoc* Dunn’s multiple comparisons test for all comparisons; Figure 1Diii). These findings provide additional evidence for both the high degree of precision involved in CRISPR/Cas9-induced NHEJ repair, and that differences in repair are dependent on individual sgRNAs, possibly based on target sequence.

Having observed that CRISPR/Cas9-induced NHEJ repair does not appear to be inherently error-prone, we next sought to examine whether a given pair of sgRNAs generates similar repair patterns in different cell types. Thus, we designed pairs of sgRNAs against the PAX7 3’UTR, which were subsequently transfected into HEK293 cells and H9 human ESCs (hESCs) (Figure 1Ei). We observed a mostly high level of precise repair in the H9 hESCs, with only one pair resulting in less than 75% precise repair (Figure 1Eii; Figure 1-figure supplement 3 for HEK293; Figure 1-figure supplement 4 for H9 hESCs). However, we observed what appeared to be a more varied level of precise repair in HEK293 cells with the same PAM pairs, ranging from 44.44% to 90.48% (Figure 1Eii). Additionally, we only observed two typical error-prone events for HEK293, one resulting from the use of PAX7r1-4 and a double event for PAX7r1-4 and PAX7r1-1 (Figure 1-figure supplement 3). For H9 hESCs, we observed two typical error-prone repair events: one for PAX7r1-2 and one for PAX7r1-3 (Figure 1-figure supplement 4). In spite of this variation, deviation from 0/0 precise repair was not significant for any given pair between cell types. In fact, for each pair, the deviation from 0/0 precise repair was not significantly different between cell types (*P* > 0.8 for all four comparisons; Dunn’s multiple comparisons test; Figure 1Eiii). These data illustrate that a given pair of sgRNAs generates similar patterns of repair with similar degrees of precision in different cell types of the same species.

Given that we observed a high degree of precise repair in human cells across three loci and 16 pairs of sgRNAs, we sought to examine if a high level of repair precision would also be observed in mouse cells. Paired sgRNAs have been previously utilized to generate deletions in murine embryos, although there was significant variability in deletion size between embryos (Zhou, Wang *et al*., 2014). To examine repair patterns in mouse cells, we transfected C2C12 murine immortalized myoblast cells with three pairs of sgRNAs in different orientations against the mUtrn 3’UTR (Figure 1Fi). We observed not only a more varied pattern of repair than with human cells (precise repair ranged from 41.67% to 100% of amplicons; Figure 1Fii; Figure 1-Figure supplement 5), but also a higher degree of repetition in the imprecise excision repair amplicons in their deviation from 0/0 precise repair (Figure 1Fiii). Interestingly, we found that there was no significant difference between the distributions of deviance from 0/0 precise repair for the three orientations (*P* = 0.1354; Kruskal-Wallis test). These results indicated that CRISPR/Cas9 facilitates a high level of repair in murine cells as well, but may potentially have generated a more varied distribution of imprecise repair due to species-specific differences in NHEJ repair.

### Knock-in Blunt Ligation: CRISPR/Cas9-Mediated in vivo Blunt-End Cloning

Because of the high level of precise NHEJ repair we observed, we asked if we could exploit this apparent *in vivo* blunt-end ligation to replace endogenous sequences precisely with linear exogenous sequences in a homology-independent manner using CRISPR/Cas9 (Figure 2A). A similar method has been developed in zebrafish using plasmid DNA in which the same sgRNA is used to cleave both the genome and the plasmid, ultimately resulting in homology-independent knock-in of the plasmid, albeit with reduced levels of perfect ligation (Auer *et al*., 2014). To test this idea and examine the level of repair precision, we constructed an expression cassette consisting of the mouse PGK promoter driving expression of the Puro^R^ΔTK fusion gene linked via a P2A skipping peptide to mCherry, followed by the rabbit beta globin terminator sequence (PTKmChR). We also constructed another cassette containing the same sequences, but flanked by a phiC31 *attP* site upstream of the PGK promoter and a Bxb1 *attP* site downstream of the terminator sequence for the initial purpose of generating a selectable landing pad that did not rely on homology arms for integration (DICE-EPv2; Figure 2B). Each cassette was amplified via PCR with 5’ phosphorylated primers, purified, and subsequently co-transfected into HEK293 cells with various sgRNA/Cas9 constructs, singly or in pairs, before being subjected to flow cytometric analysis and junction PCR, followed by Sanger sequencing (Figure 2C). In this set of experiments, we utilized sgRNAs targeting the PAX7 3’UTR (Figure 2D). For the DICE-EPv2 cassette, flow cytometric analysis revealed that the percentage of mCherry+ cells was increased above cassette alone (0.20 ± 0.15%) for sgRNAs PAX7r2-1 and PAX7r3-2, both alone and paired (referred to as PAMs Out Large) with a range 1.61-2.67% two days post-transfection, and that the percentage of mCherry+ cells could be further increased to 17.80-24.50% with four days of puromycin selection, albeit at the risk of increasing selection for random integration (Figure 2E-F; 14.30 ± 0.20% for cassette alone). For the PTKmChR cassette after two days, we also observed that the percentage of mCherry+ cells was increased above cassette alone (0.56 ± 0.32%) in the presence of all sgRNAs tested, which consisted of two pairs of PAMs 5’, two pairs of PAMs Out (the four pairs previously characterized in Figure 1F), and the PAMs Out Large pair, together and singly for a range of 2.63-5.74% (Figure 2G). Interestingly, four days of puromycin selection resulted in a lower percentage of mCherry+ cells with cassette alone (6.57 ± 1.64%) than with the DICE-EPv2 cassette. Puromycin selection also resulted in an increase in the percentage of mCherry+ cells for the PAMs 5’ and PAMs Out pairs, ranging from 17.10% to 32.60%.

**Figure 2.**
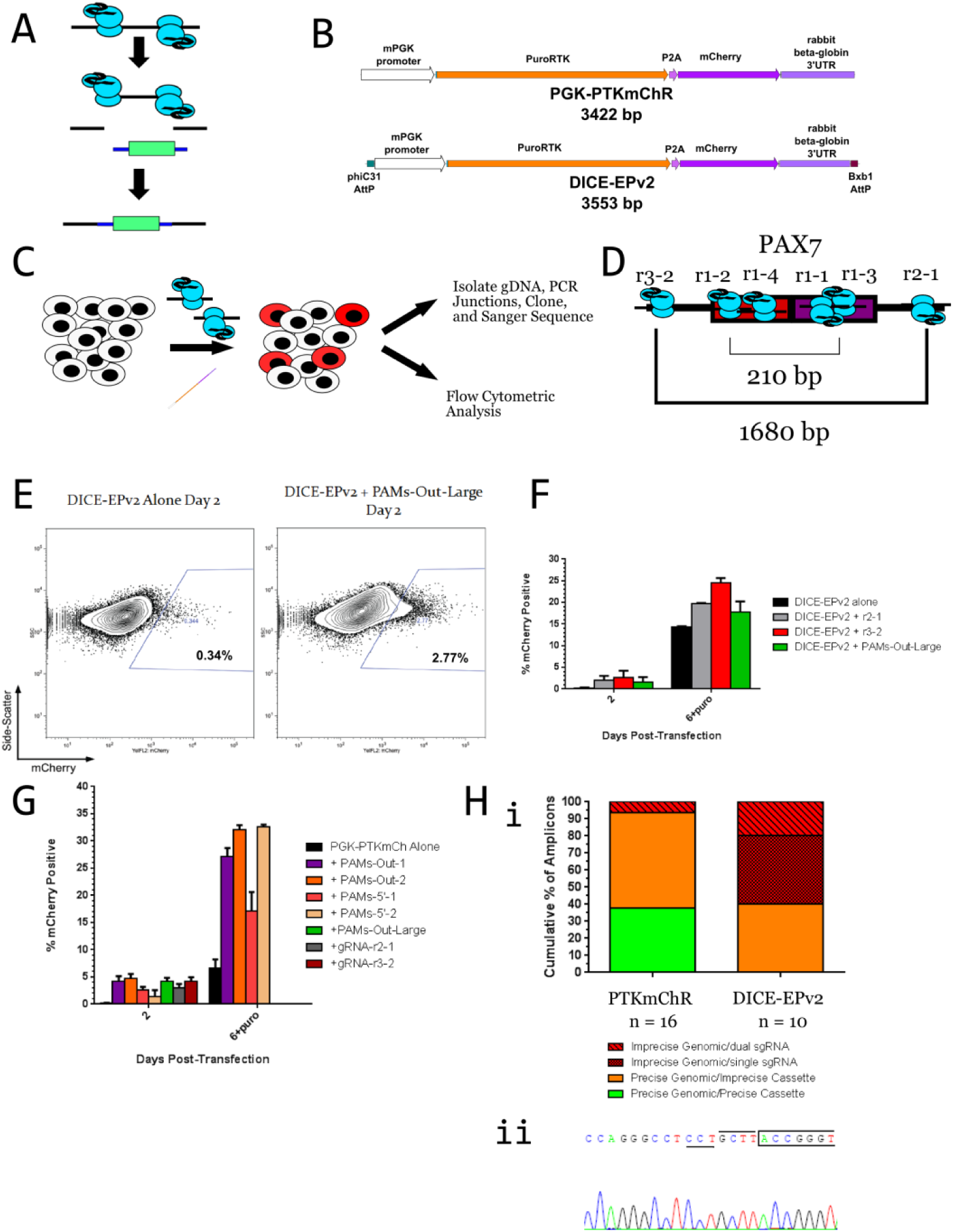
Knock-in blunt ligation (KiBL) in human immortalized cells. (A) The use of two sgRNAs at the same locus should facilitate ligation of exogenous sequence in place of the excised sequence. (B) Schematic of the PGK-PTKmChR and DICE-EPv2 cassettes. (C) Workflow for KiBL experiments: HEK293 cells are transfected with sgRNA/Cas9 vectors and cassettes before flow cytometric analysis and sequence analysis. (D) Schematic diagram of the PAX7 3’UTR region along with sgRNAs used in these experiments. (E) Representative flow cytometric data displaying the difference in the percentage of mCherry+ cells between the DICE-EPv2.0 cassette transfected alone and with sgRNA/Cas9 vectors after two days. (F) Quantification of the DICE-EPv2 transfection experiments in terms of percentage of mCherry+ cells using sgRNAs PAX7r2-1 and PAX7r3-2. *n* = at least 2 independent experiments consisting of 3 replicates each for each transfection Data is shown as the mean ± SEM. (G) Quantification of the PGK-PTKmChR cassette transfections. Same experimental conditions as in panel F. *n* = at least 2 independent experiments consisting of 3 replicates each for each transfection. Data is shown as the mean ± SEM. (H) (i) Graph of cumulative percentage of sequenced amplicons categorized by repair status of the genome-cassette junction. *n* for each is displayed on the graph. (ii) An example chromatogram of the precise genomic/precise cassette junction for the PTKmChR cassette. Boxed sequence denotes PGK promoter sequence, PAM is underlined, and the first four bases of sgRNA target are overlined.

Subsequent PCR analysis of bulk unsorted cells for all possible junctions of genomic and cassette sequences revealed that knock-in blunt ligation of the DICE-EPv2 cassette led to imprecise repair of the genomic junction (60% of sequenced amplicons) and an absence of precise cassette junction repair, whereas knock-in blunt ligation of PTKmChR cassette resulted in a high level of precise genomic junction repair (93.75%) and an appreciable degree of precise cassette junction repair (37.5%; Figure 2Hi and ii). We speculated that the lack of precise cassette junction repair with DICE-EPv2 was attributable to the formation of hairpin structures by the palindromic *AttP* sites. Such hairpins could be unfavorable to direct end-joining. Thus, these hairpins would be more subject to removal by nucleases. Additionally, we observed one instance of an insertion of part of the pX330 plasmid (similarly to that observed in Canver, Bauer *et al*., 2014). These results indicated that the blunt DSBs generated by the CRISPR/Cas9 system could be exploited in mammalian cells to knock-in PCR cassettes in a homology-independent manner. We observed that cassettes were knocked into the genome in both orientations, indicating that ligation by precise NHEJ lacks directionality, as expected due to lack of homology arms on our cassettes (Figure 2-figure supplement 1).

Because of the relatively large size of the DICE-EPv2 and PTKmChR cassettes and the relative dimness of mCherry, we developed a second series of smaller, brighter cassettes. These cassettes consist of the Ef1α promoter driving the expression of a fluorescent protein, which is followed by the rabbit β-globin terminator sequence (Figure 3A). Our fluorescent proteins of choice were Clover, mRuby2 (both from Lam *et al*., 2012), mKOκ (Tsutsui *et al*., 2008), mCardinal (Chu *et al*., 2014), and mCerulean (Rizzo *et al*., 2004) due to minimal spectral overlap and (for the most part) brightness. These cassettes were less than 2 kb in size and could be readily amplified from template plasmid and easily purified. We called these cassettes the pKER (polymerase-chain-reaction-based <u>K</u>nock-in <u>E</u>F1α-<u>R</u>BG terminator) series, and they are depicted by brightness in Figure 3A.

**Figure 3.**
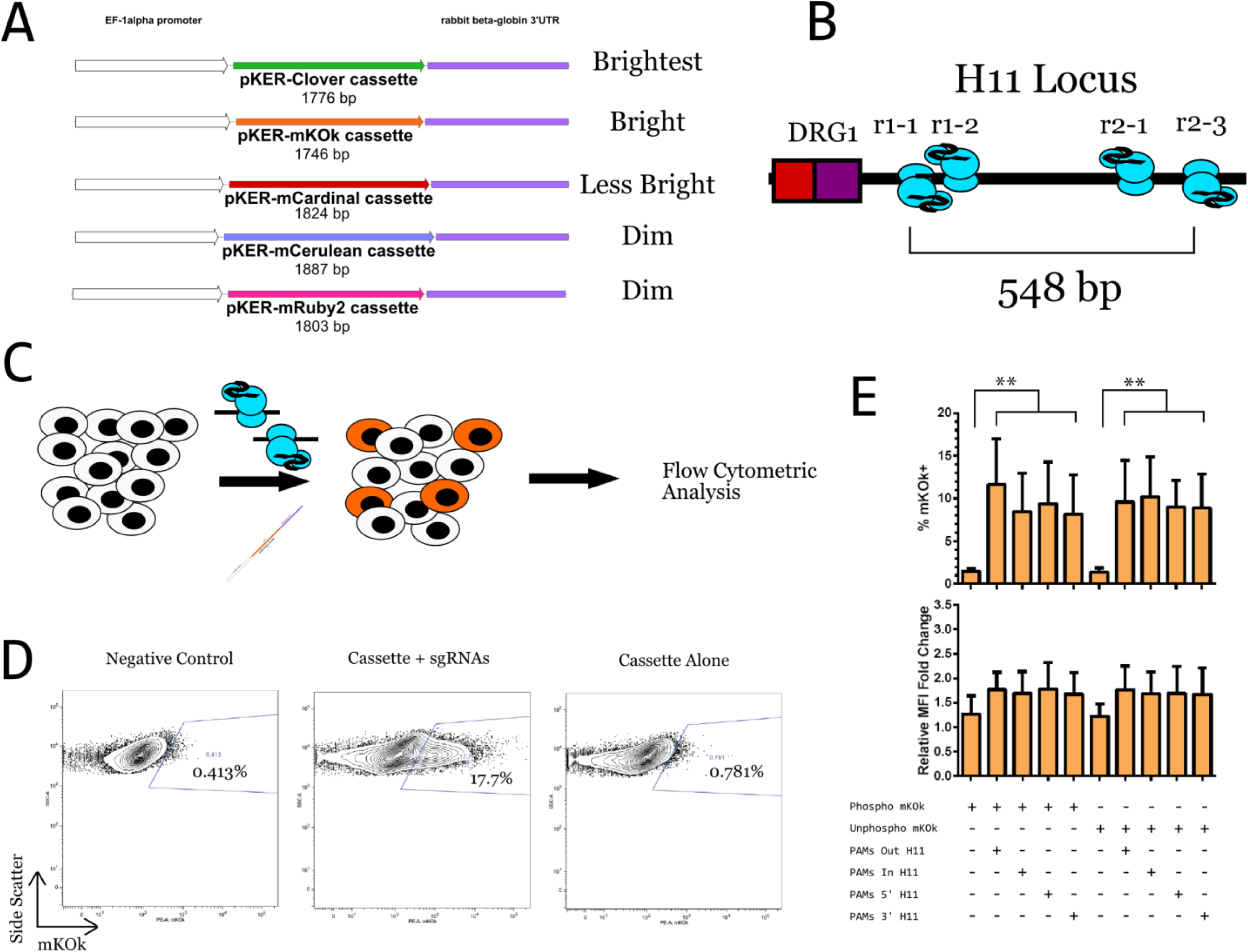
pKER cassettes are suitable for use in knock-in blunt ligation. (A) Schematic diagrams of the pKER series of cassettes organized by brightness. Each cassette consists of a fluorescent protein under the control of the EF1α promoter and followed by the rabbit beta-globin terminator sequence. (B) Schematic diagram of the 5’ side of the H11 locus on human chromosome 22 depicting the four sgRNAs positioned relative to the sense strand. 548 bp separates the 5’-most and the 3’-most sgRNAs from each other. (C) Diagram depicting the workflow for analyzing pKER expression. pKER cassettes and pairs of sgRNA/Cas9 vectors are co-transfected into HEK293 cells and analyzed by flow cytometry after two days. (D) Representative flow cytometric data generated with pKER-mKOκ and the H11 sgRNAs. (E) Quantification of the percentage of mKOκ+ cells and their relative median fluorescence intensity for phosphorylated and unphosphorylated cassettes for all four possible PAM orientations. *n* = 3 independent experiments each consisting of 3 technical replicates. Data is displayed as the mean ± SEM of the averages of each experiment and were analyzed using a two-way ANOVA with repeated measures followed by a *post hoc* Tukey’s multiple comparisons test. ** = *P* < 0.003.

To quickly assess the suitability of the pKER series for KiBL, we used pKER-mKOκ to investigate the effects of the orientation of the paired PAMs as well as the effects of the cassette’s phosphorylation status on the efficiency of KiBL, using the percentage of positive cells as a proxy. We reasoned that unphosphorylated cassettes may have a selective advantage for KiBL as they may not be detected as free, broken ends of DNA as readily as phosphorylated cassettes may be, and they may be decrease the possibility of amplicon concatemerization. For these experiments, the sgRNA pairs consisted of all four possible PAM orientations directed against the same region of the human H11 locus on chromosome 22 (Figure 3B), which was previously identified as a safe harbor site in the human genome (Zhu *et al*., 2014). H11 is an intergenic locus between *DRG1* and *EIF4ENIF1* on human chromosome 22 (Figure 3-figure supplement 1). There is a potential non-coding RNA at this locus, but it has only been observed in human lung and only from a cDNA screen. We transfected HEK293 cells with phosphorylated or unphosphorylated pKER-mKOκ cassettes and pairs of sgRNA/Cas9 vectors, before subjecting the cells to flow cytometric analysis two days later (Figure 3C). Two days post-transfection, we observed a higher percentage of mKOκ+ cells in the cultures co-transfected with CRISPR/Cas9 vectors (ranging from 8.15-11.64%) than in those transfected with cassette alone (1.34-1.49%; Figure 3D). Surprisingly, flow cytometry did not reveal any significant differences between PAM orientations or cassette phosphorylation status in terms of percentage of mKOκ+ cells or relative normalized median fluorescence intensity (MFI), other than that the presence of Cas9 increased both measures (Figure 3E; two-way ANOVA with repeated measures followed by post-hoc Tukey’s multiple comparisons test, P < 0.003 for all comparisons when comparing PAMs-treated to cassette alone). These results suggested that the pKER cassettes were well-suited for use in KiBL: positive cells were bright, and treatment with Cas9 resulted in a large increase in positive cells.

### Stabilized Cas9-DD Combined with Nuclease Protection Facilitates a Higher Degree of Precise Repair at the Genome-Cassette Junction

We and others have observed that, in utilizing the CRISPR/Cas9 system, successful modifications of one allele appeared to coincide with undesired mutations in the sister allele (unpublished data; Canver, Bauer *et* al., 2014). The duration of Cas9 activity may be responsible for these additional modifications. Thus, we initially sought to generate a more temporally controllable Cas9. We chose to utilize the FKBP12 L106P destabilization domain because it was relatively small, has been shown to be effective at destabilizing a wide range of proteins, and its stabilizing ligand, the small molecule Shield-1, was relatively inexpensive and had virtually no effect on cell viability, as well as good intracellular availability and potency (Banaszynski *et al*., 2006; Maynard-Smith *et al*., 2007). We initially chose to fuse the destabilization domain to the C-terminus of SpCas9 because we desired a version of Cas9 that possessed some residual stability in the absence of Shield-1 (Banaszynski *et al*., 2006). The resulting fused domain was in close proximity to the PAM-binding domain of SpCas9 (Anders *et al*., 2014). We called this version of Cas9 “Cas9-DD” (Figure 4A, Figure 4-figure supplement 1A). Additionally, we also constructed a version of Cas9 with the destabilization domain fused to the N-terminus, which we called “DD-Cas9” (Figure 4-figure supplement 1A-B). We then examined the relative stability of Cas9-DD and DD-Cas9 via Western blotting of HEK293 cells transfected with constructs encoding the H11-r1-2 sgRNA and WT-Cas9, Cas9-DD, or DD-Cas9 (Figure 4-Figure supplement 1C). After 29 hours, we observed a higher level of Cas9-DD protein present in cells treated with 0.5 μM Shield-1, compared to cells treated with Cas9-DD vector alone (destabilized Cas9-DD). Similarly, for DD-Cas9, treatment with 0.5 μM Shield-1 appeared to increase the level of DD-Cas9 protein relative to DD-Cas9 vector alone. These results indicated that the addition of the destabilization domain to Cas9 resulted in a destabilized protein that could be stabilized by the addition of Shield-1.

**Figure 4.**
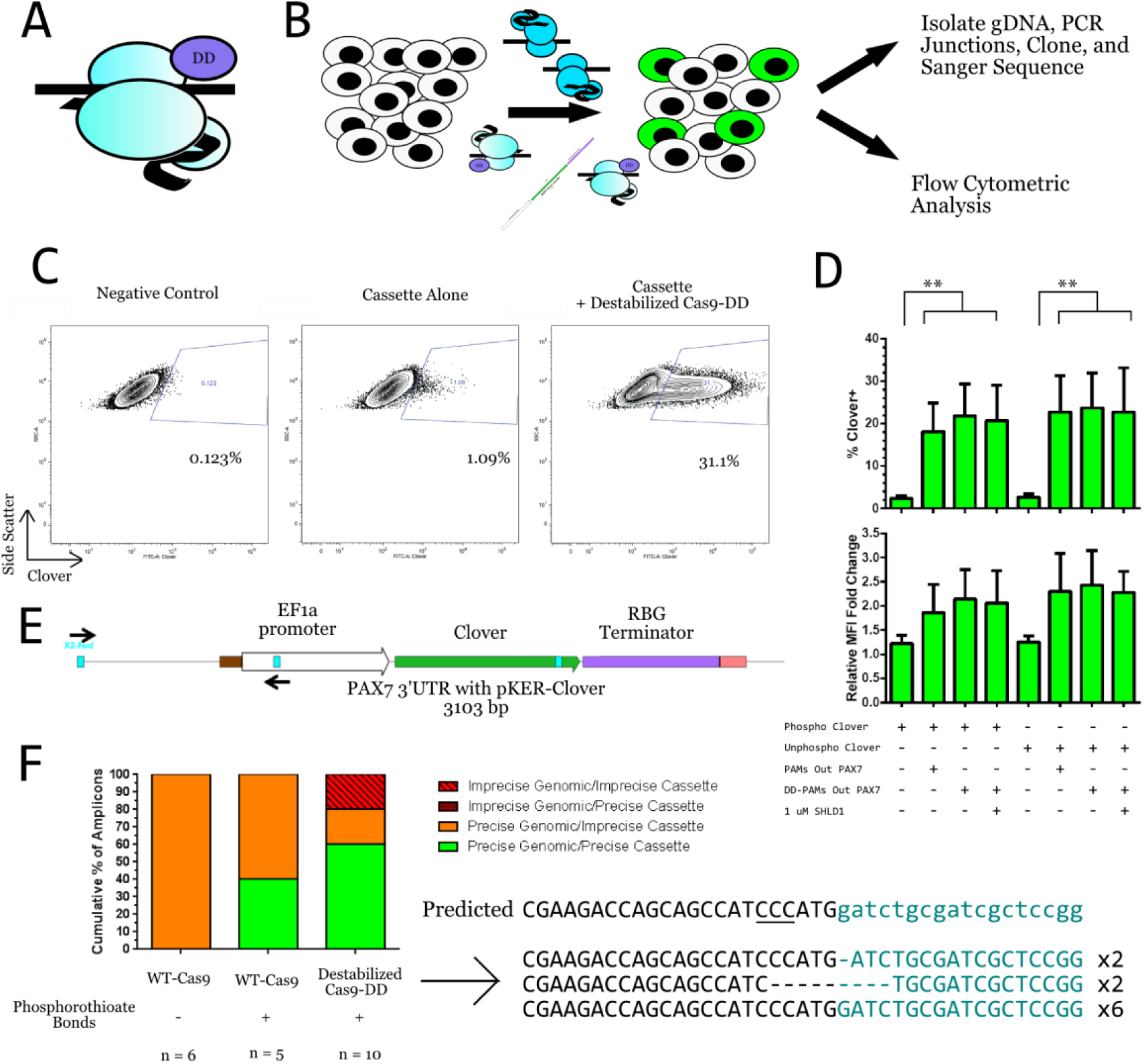
Cas9-DD and nuclease protection facilitate a higher degree of precise genome-cassette junction repair. (A) Diagram of Cas9-DD showing the location of the destabilization domain on SpCas9. The domain is near the PAM-binding domain at the C-terminus. (B) Schematic diagram displaying the workflow for comparing WT-Cas9 to Cas9-DD. HEK293 cells are transfected with pKER-Clover cassette and the PAX7 PAMs Out-Large pair of vectors encoding either WT-Cas9 or Cas9-DD. Two days post-transfection, a portion of the cells are subjected to flow cytometric analysis and the remainder are reserved for genomic DNA isolation for sequence analysis. (C) Representative flow cytometric data for the pKER-Clover cassette and the PAX7 PAMs Out-Large sgRNAs with destabilized Cas9-DD. (D) Quantification of the percentage of Clover+ cells and their relative median fluorescence intensity for phosphorylated and unphosphorylated cassettes for WT-Cas9, destabilized Cas9-DD, and stabilized Cas9-DD (1 μM Shield-1 for 24 hours). *n* = 5 independent experiments each consisting of 3 technical replicates. Data is displayed as the mean ± SEM of the averages of each experiment and were analyzed using a two-way ANOVA with repeated measures followed by a *post hoc* Tukey’s multiple comparisons test. ** = *P* < 0.001. (E) Diagram of the modified PAX7 3’UTR locus. This diagram displays pKER-Clover placed at the locus in the 5’ to 3’ orientation. Arrows indicate the primers used to generate amplicons for the sequence analysis in (F). (F) Analysis of genome-cassette junctions for the comparison of WT-Cas9 with destabilized Cas9-DD and the determination of the effect of protecting the cassette from nuclease degradation. Numbers of amplicons analyzed appear beneath their respective treatments. Also depicted are the sequences of the amplicons from the destabilized Cas9-DD junction analysis. Black denotes genomic sequence, turquoise denotes cassette sequence, PAM is underlined, and dashes denote unobserved sequence.

To characterize how Cas9-DD would affect KiBL efficiency, we transfected our PAX7 PAMs Out Large sgRNA pairs on vectors encoding the sgRNAs and either wild-type Cas9 (WT-Cas9) or Cas9-DD, along with phosphorylated or unphosphorylated pKER-Clover cassettes into HEK293 cells, which were subjected to flow cytometric analysis and harvested for genomic DNA after two days (Figure 4B). We chose PAX7 because we had already successfully carried out KiBL and analyzed the pattern of repair at this locus (Figure 2). Flow cytometric analysis confirmed that co-transfection with the sgRNA/Cas9 vectors increased the percentage of Clover+ cells (18.12-35.8%) over that of cassette alone (2.36-2.64%; Figure 4C-D; *P* < 0.002 for all comparisons; Tukey’s multiple comparisons test). Interestingly, we did not detect any significant difference between Cas9-DD and WT-Cas9 for the percentage of Clover+ cells or for normalized MFI (Figure 4D). Additionally, we did not detect any significant difference between the presence and absence of 1 μM Shield-1 with Cas9-DD.

Having already observed that WT-Cas9 facilitated knock-in blunt ligation of linear cassettes, we next sought to determine the effect of Cas9-DD at the molecular level. To this end, we examined the 5’ PAX7 genomic-5’ pKER-Clover junction resulting from KiBL using unphosphorylated cassettes with WT-Cas9 or destabilized Cas9-DD with our sgRNAs (Figure 4E, Figure 4-figure supplement 2). We did not examine junctions using stabilized Cas9-DD at this time as we reasoned that its junctions would resemble those of WT-Cas9. Additionally, we examined whether the addition of three phosphorothioate bonds in the 5’ ends of the primers used to amplify the cassettes affected the precision of the cassette side of the junction by protecting the cassette from nuclease degradation. We observed that unprotected cassettes co-transfected with WT-Cas9 resulted in precise genomic junctions, but imprecise cassette junctions in 100% of observed amplicons (Figure 4F). Interestingly, protecting the 5’ ends of the cassette only resulted in a relatively modest increase in precise genomic/precise cassette junctions (40% of amplicons) when using WT-Cas9 (Figure 4F). When destabilized Cas9-DD was used in conjunction with phosphorothioate bonds, we observed a greater increase in precise genomic/precise cassette junctions (60% of amplicons), but, surprisingly, we also observed a slight increase in imprecise genomic/imprecise cassette junctions (20% of amplicons; Figure 4F). These results appeared to demonstrate that the use of destabilized Cas9-DD, along with the addition of nuclease protection to the pKER cassettes, facilitated a higher degree of cassette junction precision in KiBL.

### KiBL results in efficient, precise knock-ins in human induced pluripotent stem cells

Having observed that the CRISPR/Cas9 system could facilitate the precise knock-in of cassettes in an immortalized cell line, we sought to apply KiBL to human iPSCs (hiPSCs). The successful application of such a method would provide a viable alternative to traditional nuclease-mediated homologous recombination (HR). We chose a line of hiPSCs generated from a healthy donor that we refer to as JF10, and chose H11 as our target locus utilizing the PAMs Out sgRNAs. The JF10 cells fortuitously possessed two allele-specific SNPs, one flanking each sgRNA target, which allowed us to determine which allele was targeted for genome editing (Figure 5A). In order to minimize the effect of untransfected cells on downstream analyses, we isolated fluorescent-protein+ cells via FACS four to seven days post-electroporation with unphosphorylated pKER cassettes containing phosphorothioate bonds and the sgRNA/Cas9 vector pairs (Figure 5B). To our surprise, we observed rather low cell viability and relatively low numbers of Clover+ cells when using the WT-Cas9 vectors, whereas we observed greater cell viability and higher numbers of Clover+ cells when using the Cas9-DD vectors in the absence of Shield-1 (Figure 5C). 0.5 μM and 1 μM of Shield-1 resulted in a moderately increased level of cell viability compared with WT-Cas9. FACS analysis revealed that the Clover+ cells formed a mostly discrete population in these hiPSCs (Figure 5D). Additionally, the percentage of Clover+ cells was always observed to be less than 0.5% of live cells (Figure 5E). This low percentage may be due both to the relatively low amount of transfected cassette (200ng) and the lower electroporation efficiency of linear double-stranded DNA. However, we found that there was a significant difference between the mean percentages of Clover+ cells (*P* = 0.0193; ordinary one-way ANOVA), but that only the mean percentages of cassette alone (0.069%) and Cas9-DD + 0.5 μM Shield-1 (0.147%) were significantly different from each other (*P* = 0.019; *post hoc* Tukey’s multiple comparisons test). In order to analyze precise end-joining at the junctions, we carried out targeted genomic amplification on pooled Clover+ cells by amplifying across the H11 locus in order to capture all of the KiBL events (Figure 5F). Subsequent analysis of the H11 5’-Clover 5’ junction revealed that, in the presence of Shield-1, Cas9-DD-induced DSBs resulted in the precise joining of both the genomic and cassette sequences, but that the absence of Shield-1 resulted in the loss of the PAM and the three bases preceding it on the genomic side, but no loss of bases on the cassette side (Figure 5G). Additionally, the use of destabilized Cas9-DD appeared to result in a C-to-T mutation nine bases 5’ of the PAM. We have only observed this mutation with Cas9-DD in the absence of Shield-1. Bulk sequencing of the Clover 3’-H11 3’ junction for stabilized Cas9-DD also showed precise joining of the genome with the cassette (Figure 5-figure supplement 1A). When utilizing stabilized DD-Cas9, we observed precise genomic repair and mostly precise cassette repair, but we also observed the same genomic deletion and C-to-T mutation at the 5’ junction as we had observed with destabilized Cas9-DD (Figure 5-Figure supplement 1B). When analyzing destabilized DD-Cas9, we did not observe the presence of 5’ H11-5’cassette junctions (Figure 5-Figure supplement 2). These results demonstrated that KiBL is feasible in hiPSCs and that Cas9-DD may be preferable to WT-Cas9 due to increased cell viability. Additionally, these results demonstrated a difference between the stabilized and destabilized forms of Cas9-DD in that stabilized Cas9-DD may result in higher precision KiBL whereas destabilized Cas9-DD may be more useful for probing repair dynamics of Cas9-induced DSBs.

**Figure 5.**
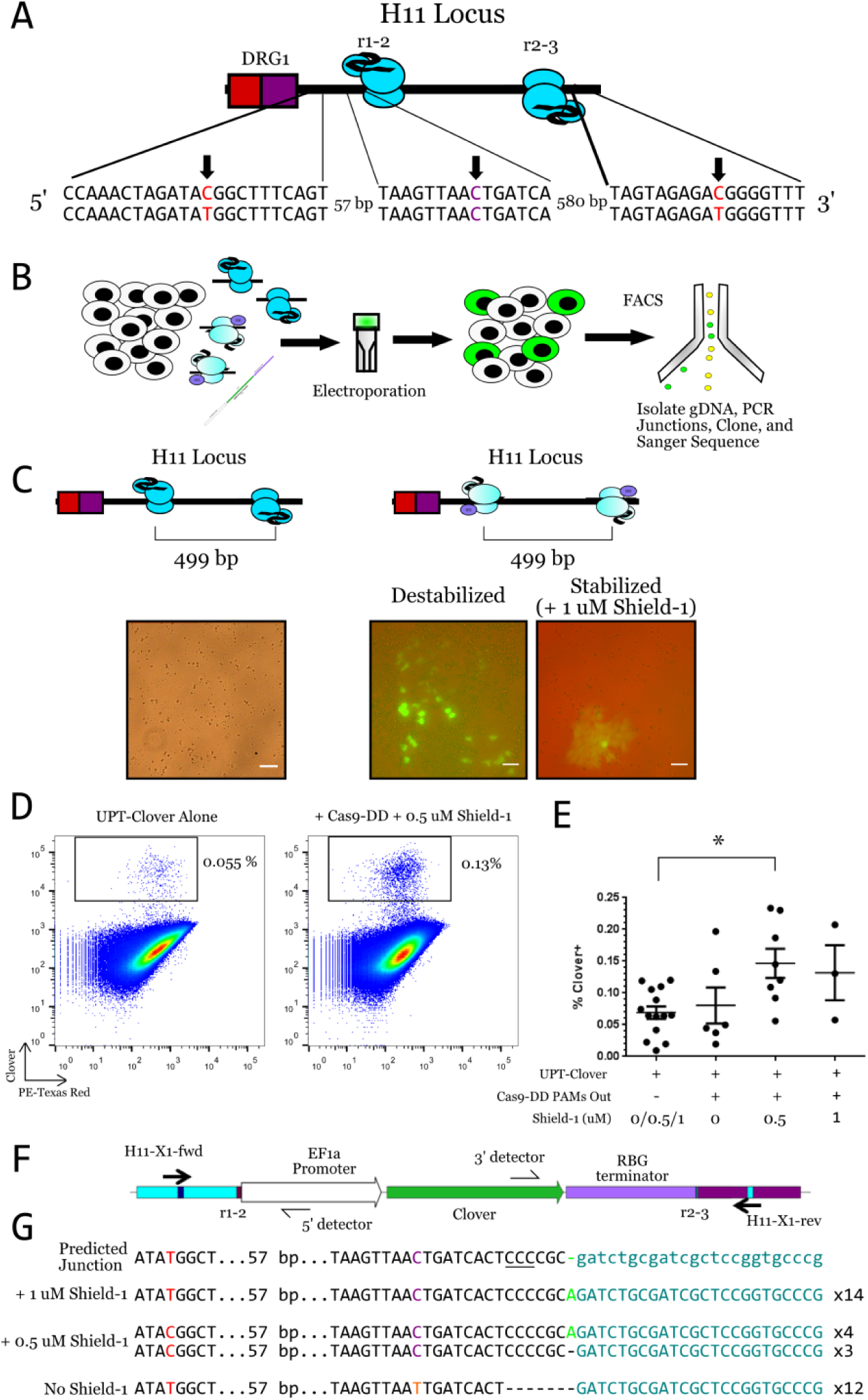
Stabilized Cas9-DD facilitates knock-in blunt ligation in hiPSCs. (A) Schematic diagram of the targeted H11 locus in JF10 hiPSCs showing the sequence variation at three regions. Individual bases of interest are indicated with an arrow. SNPs are indicated in red. The middle region depicts a homozygous cytosine in violet 9bp upstream of the PAM of the 5’ sgRNA that differs from human reference sequence. (B) Diagram of the workflow for examining the efficiency KiBL in hiPSCs. Cells are electroporated with 200 ng of unphosphorylated phosphorothioate (UPT) pKER-Clover and 500 ng of each vector encoding the H11 PAMs Out sgRNA pair for WT-Cas9 or Cas9-DD. Four to seven days later, Clover+ cells are isolated with FACS for genomic DNA extraction and subsequent downstream analysis of genome cassette junctions. (C) Comparison of WT-Cas9- and Cas9-DD-mediated KiBL at the H11 locus via fluorescence microscopy. Scale bar = 50 μm. (D) Examples of FACS plots from KiBL using UPT-Clover. Plots are of singlet live cells. (E) Quantification of the percentage of Clover+ cells via FACS for KiBL using UPT-Clover and Cas9-DD. *n* for each condition = at least 2 independent experiments with at least 2 technical replicates each. Each data point represents the mean percentage of Clover+ cells for one experiment. The mean of the experiments is graphed as a horizontal line; the error bars are ± SEM. Data were analyzed using an ordinary one-way ANOVA with a post-hoc Tukey’s multiple comparisons test. * = *P* < 0.02. (F) Schematic diagram showing targeted amplification strategy of knock-in blunt ligation events at the H11 locus. Full arrowheads denote the primers used to amplify across the locus in the primary PCR. Half arrowheads denote the primers used in conjunction with the full arrowheads to amplify the genome-cassette junctions via nested PCR. (G) Sequence analysis of the 5’-genome 5’-cassette junctions of the KiBL event depicted in panel (F) for various degrees of stability of Cas9-DD. 5’ allele-specific SNP is denoted in red. The 5’ side of the pKER cassette is denoted in turquoise. Nucleotide in green corresponds to Cas9 cleaving at the fourth base upstream of the PAM (underlined) instead of the third. The cytosine colored violet indicates the reference base for JF10 at that position; the thymine in orange indicates a mutation not present in JF10. Dashes indicate unobserved bases.

We recognized that there was the possibility that we were repeatedly sampling and sequencing the same clonal population in our initial hiPSC KiBL experiments. We attempted to rule out this possibility in three ways. First, we carried out single-cell PCR across the H11 locus utilizing FACS to sort single cells into wells of 96-well plates. From this experiment, we identified two cells that underwent KiBL: one originated from treatment with stabilized Cas9-DD (cell I), the other from destabilized Cas9-DD (cell II) (Figure 6A). Cell I, interestingly, appeared to have only received the 5’ sgRNA/Cas9-DD vector and had precise genomic and cassette junctions for both the 5’-5’ and 3’-3’ junctions. Sequence analysis revealed that cell II received both sgRNA vectors and possessed a precise 5’ genomic junction, but was lacking four bases of the 5’ cassette junction. Interestingly, the 3’ cassette junction was precise, but the 3’ genomic junction appeared to have been cut 16 bases downstream of the actual PAM site for the 3’ sgRNA (Figure 6A). In both cells, the other allele appeared be completely unmodified except for an additional adenosine present at the cleavage site of the 5’ sgRNA, potentially as the result of error-prone NHEJ.

**Figure 6.**
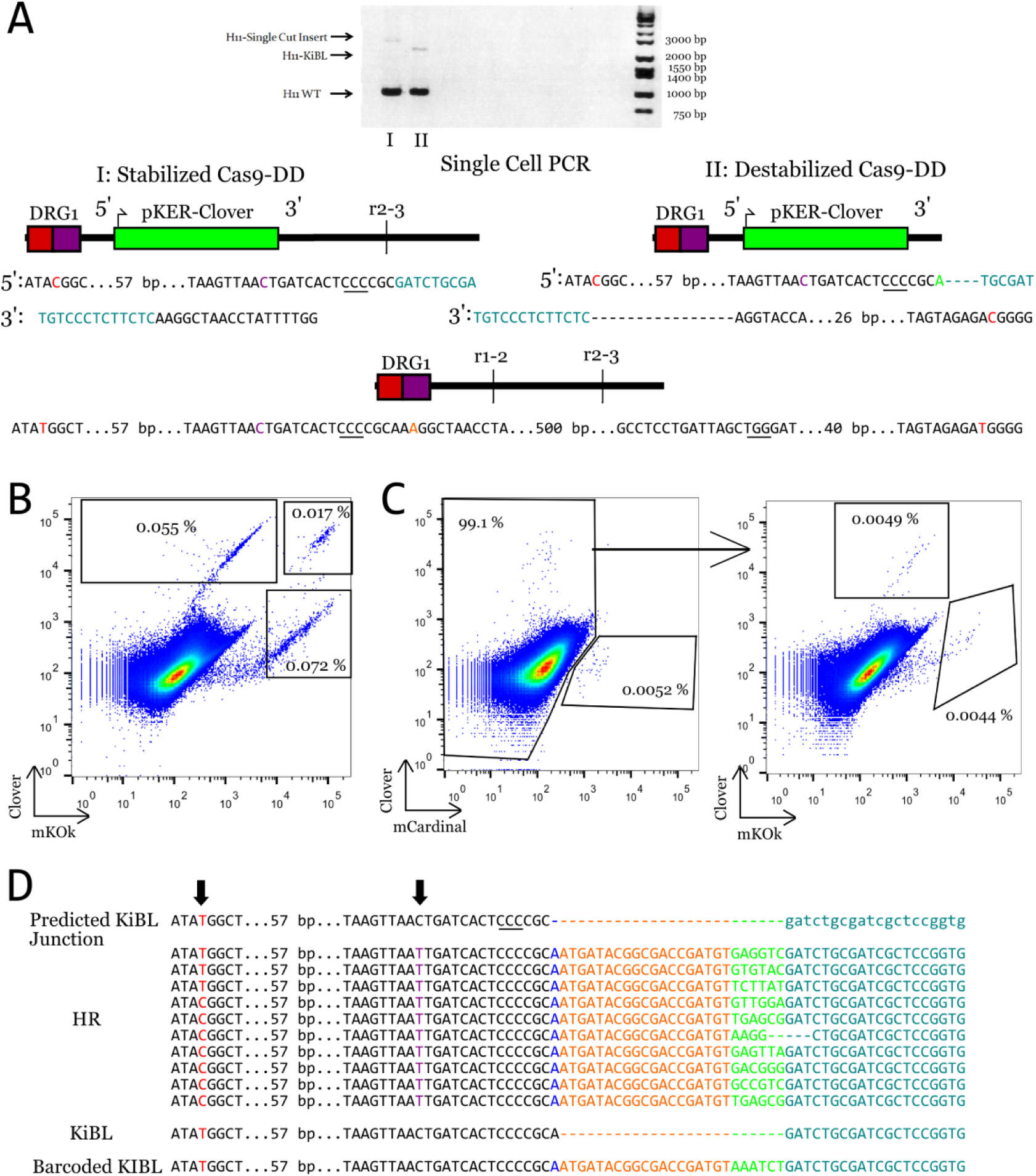
Clonality analysis of KiBL using hiPSCs at the H11 locus. (A) Single-cell PCR-based analysis of two cells from independent transfections. Red denotes allele-specific SNPs and *de novo* mutations, turquoise denotes cassette sequence, black denotes genomic, green denotes wobble-cleaved bases, violet denotes the wild type SNP for JF10 that differs from reference, dashes denote unobserved sequence, and orange denotes an insertion. (B) Representative FACS plot of singlet live hiPSCs transfected with equal amounts (100 ng each) UPT-Clover and UPT-mKOκ cassettes with stabilized Cas9-DD the H11 PAMs Out sgRNA pair. Data is representative of two independent experiments. (C) FACS plot of singlet live hiPSCs transfected with equal amounts (66.67 ng each) UPT-Clover, -mKOκ, and mCardinal cassettes with stabilized Cas9-DD the H11 PAMs Out sgRNA pair. (D) Sequence analysis KiBL events at 5’ H11-5’ pKER junction resulting from the use of stabilized Cas9-DD and barcoded UPT-Clover cassettes possessing short 50 bp homology arms. Red denotes allele-specifc SNPs (also indicated by an arrow), violet denotes HR-mediated conversion of a base (also indicated by an arrow), blue denotes a base HR-mediated insertion, orange denotes a common index sequence, green denotes the barcode sequence, turquoise denotes the 5’ cassette sequence, dashes indicated unobserved sequence. PAM is underlined.

To further examine the level of clonality generated by KiBL, we took advantage of the pKER cassettes’ limited spectral overlap and co-transfected 1) pKER-Clover and pKER-mKOκ, and 2) pKER-Clover, -mKOκ, and –mCardinal with our H11 PAMs Out sgRNA pair into JF10 hiPSCs, which were subsequently analyzed via FACS. In the first condition, we observed that Clover+mKOκ- and Clover-mKOκ + cells were roughly equivalent in relative frequency and that Clover+mKOk + cells made up a smaller frequency of the population (Figure 6B). Interestingly, in the second condition, we observed all three possible single-positive populations, and no double- or triple-positive populations (Figure 6C). These data illustrated that there were at least three clones in each experiment.

To address clonality at the molecular level, we employed an approach using pKER-Clover cassettes containing 6-base barcodes and 50-base homology arms. Homology arms of this size have been previously used with the CRISPR/Cas9 system to facilitate homologous recombination in human immortalized cell lines (Zheng, Cai *et al*., 2014) and mouse ESCs (Li *et al*., 2014). After co-transfection with the H11 PAMs Out sgRNA pairs and Cas9-DD, followed by isolation with FACS, sequence analysis of the 5’ genomic-5’ Clover junction revealed that HR occurred several times (Figure 6D). After isolation with FACS, PCR amplification of the 5’ genomic-5’ Clover junction, and subsequent subcloning, we observed 11 different barcodes in 12 sequenced amplicons (Figure 6D). Out of these, 11 had a thymine (the human reference allele, which we had placed in the 5’ homology arm) instead of cytosine (the allele possessed by JF10) in the position prior to the PAM that differed from reference, indicating that homologous recombination did indeed occur. We also observed that both alleles underwent HR, as indicated by detection of both alleles of our allele-specific T/C SNP on the 5’ side of the locus. Additionally, we observed one non-barcoded KiBL event and a barcoded KiBL event where the cassette appeared to lack at least the 5’ homology arm. We did not observe any KiBL events where the homology arm was incorporated via NHEJ rather than HR. These results indicated that several independent KiBL events occurred in the Clover+ populations, ruling out concerns over clonal amplification affecting our analyses. Additionally, these data demonstrated that PCR amplicons possessing short homology arms combined with the CRISPR/Cas9 system facilitated homologous recombination in hiPSCs, which had not previously been demonstrated.

### Analysis of Off-Target KiBL Events

Because KiBL utilizes the CRISPR/Cas9 system, there was a concern that off-target cleavage would result in the uptake of cassettes at sites other than the targeted locus. Indeed, such events have been observed in HEK293T cells using the GUIDE-seq method for unbiased identification of CRISPR/Cas9 off-target cleavage events (Tsai *et al*., 2014b). We chose to examine the top six off-target sites for each sgRNA in the H11 PAMs Out pair as determined by the MIT CRISPR Design tool. For each off-target site, we attempted to amplify the 5’ genomic-5’ cassette, 3’ genomic-5’ cassette, 5’ genomic-3’ cassette, and 3’genomic-3’ cassette junctions, which led to 48 off-target reactions per sample. In order to perform such a large number of assays, we used the multiple displacement amplification variation of whole genome amplification to ensure that there would be enough DNA. We analyzed seven pools of Clover+ cells: one pool treated with WT-Cas9, two pools treated with destabilized Cas9-DD, two pools treated with Cas9-DD and 0.5 μM Shield-1, and two pools treated with Cas9-DD and 1 μM Shield-1. We were unable to detect off-target KiBL events at the vast majority of predicted junctions, with the exception of faint detection of the 3’H11 sgRNA OT4 genomic reverse primer-3’pKER cassette junction (Figure 7). These data suggest that, at least in hiPSCs, KiBL occurred predominantly at the on-target locus. It is also possible that the frequency of KiBL off-target events fell below the sensitivity of our assay.

**Figure 7.**
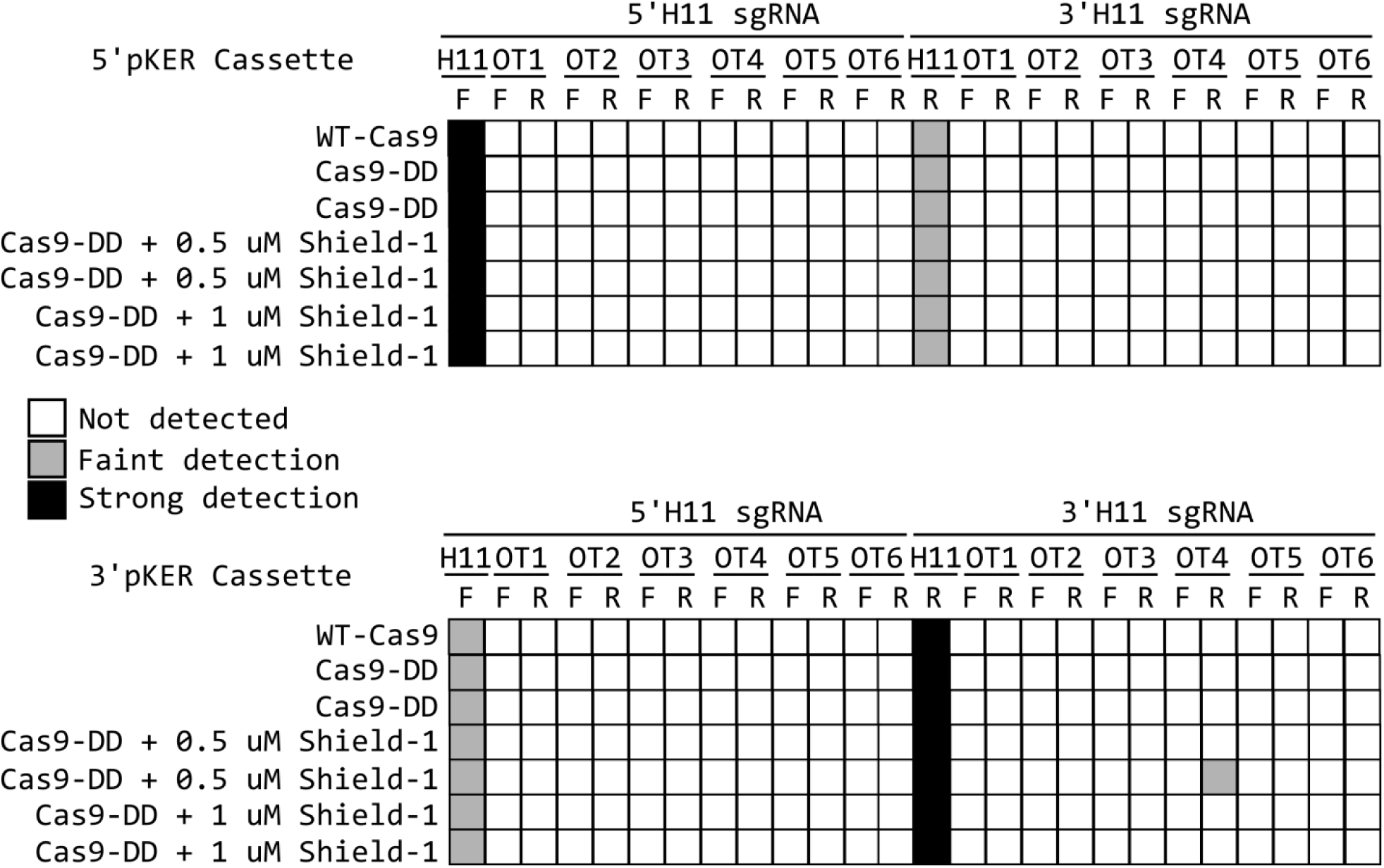
Targeted analysis of off-target KiBL events in hiPSCs. Genomic DNA from various CRISPR/Cas9-treated UPT-Clover+ cells was subjected to whole genome amplification and used for analysis of the top six off-target sites (denoted OT1-6) for both the sgRNAs of the H11 PAMs Out pair. Each off-target site was analyzed using a sense (denoted F) and an antisense (denoted R) primer designed to amplify the off-target locus. Each F and R primer were used in two amplifications, one with the 5’ pKER cassette detection primer and the other with the 3’ pKER cassette detection primer. H11 F and H11 R correspond to the primers H11-X1-Fwd and H11-X1-Rev respectively and act as on-target controls. White indicates lack of detection, grey indicates faint detection, and black indicates strong detection. Each Cas9 treatment indicates an independent pool of cells.

## Discussion

We demonstrated in this work that the blunt cleavage activity of the CRISPR/Cas9 system mediates repair with a high degree of precision and can be exploited to precisely insert exogenous sequences into the genome through knock-in blunt ligation. We found that all four paired PAM orientations are capable of facilitating precise repair and that precise repair can be observed in both human and mouse cells. We then demonstrated that PCR amplicons lacking homology arms can be used to precisely replace the sequence between sgRNA targets and that this *in vivo* blunt-end cloning functions in an immortalized cell line and in hiPSCs. Lastly, we developed a variant of Cas9 fused to a destabilization domain with the initial goal of facilitating a greater degree of control over Cas9 activity.

Other strategies utilizing the CRISPR/Cas9 system to facilitate knock-in of exogenous sequences into the genome have relied on knocking in plasmid DNA through NHEJ (Auer *et al*., 2014) or microhomology-mediated end-joining (Nakade *et al*., 2014). In general, the plasmid-based methods rely on *in vivo* cleavage of a plasmid donor to mediate integration, which was previously demonstrated with ZFNs (Cristea *et al*., 2012). These methods introduce additional undesired sequence into the cell through the plasmid backbone, whereas KiBL only introduces the knock-in cassette and the sgRNA/Cas9 vector. Additionally, the Cas9 nickase has been used to knock-in a double-stranded oligonucleotide (dsODN) with overhangs in a strategy similar to the ObLiGaRe method used with ZFNs, albeit with relatively low efficiency (Ran, Hsu *et al*., 2014; Maresca *et al*., 2013). Our observation that unphosphorylated cassettes are capable of ligating into CRISPR/Cas9-induced DSBs at relatively high frequency in HEK293 cells appears to contradict observations in the GUIDE-seq method (Tsai *et al*., 2014b). However, this finding can be explained by our much larger cassettes: our unprotected, unphosphorylated cassettes may be much less negatively affected by endogenous nucleases than the 34-bp GUIDE-seq oligos.

Another set of methods for knock-ins has been developed around exploiting HR and HDR. In cultured *Drosophila* cells, the use of PCR amplicons containing homology arms has been used to great effect (Böttcher *et al*., 2014), which is similar to what has been observed with human immortalized cell lines and mouse ESCs (Zheng *et al*., 2014; Li *et al*., 2014). A subset of these methods has focused on increasing HDR frequency through cell cycle synchronization (Lin, Staahl *et al*., 2014) and by inhibition of the NHEJ machinery (Chu *et al*., 2015; Maruyama *et al*, 2015). Our method offers the advantage of functioning during the G1, S, and G2 phases of the cell cycle, whereas the HR and HDR strategies are restricted to late S and G2 phases. Thus, our method may be particularly useful in non-dividing cells, such as neurons where the application of Cas9 has been limited to generating mutations through error-prone repair (Swiech, Heidenreich *et al*., 2014).

Targeted genome engineering has the potential to be a useful tool for research, therapeutic, and industrial applications. The CRISPR/Cas9 system has the advantages of being relatively inexpensive and easy to use, but may have larger off-target effects than TALENs or ZFNs. Several approaches have been put forward to restrict off-target cleavage, including simply using less sgRNA and Cas9, using truncated sgRNAs for increased specificity, using paired nickases, controlling Cas9 expression through the inducible Tet-ON system, and splitting Cas9 in half (Hsu *et al*., 2013; Fu *et al*., 2013; Ran, Hsu *et al*., 2013; Zhu, Gonzalez, and Huangfu, 2014; Wright, Sternberg *et al*., 2015; Zetsche, Volz, and Zhang, 2015). Tet-ON induction, however, generates a large amount of mRNA, which could lead to an increase in Cas9 off-target activity. The split-Cas9 variants, while potentially being more controllable, have substantially reduced cleavage activity, relying on either the sgRNA or rapamycin to bind the two halves together (Wright, Sternberg *et al*., 2015; Zetsche, Volz, and Zhang, 2015). Our destabilized Cas9 retains the simplicity of requiring only two expression cassettes, rather than three or four. Additionally, Shield-1 itself does not induce any undesired responses when applied to cells in culture or in animals, unlike rapamycin, (Banaszynski *et al*., 2006; Maynard-Smith *et al*., 2007; Banaszynski *et al*., 2008). During the preparation of this manuscript, a fifth variant was developed using an evolved ligand-dependent intein to restrict the activity of Cas9 until the binding of the ligand (in the form of 4-hydroxytamoxifen) results in the cleavage of the intein and activation of Cas9 (Davis *et al*., 2015). Such a system is elegant, but possesses the drawback of being unable to restrict Cas9’s activity once the intein is cleaved. It would be interesting to combine the destabilization domain with this intein because such a combination could allow for an unprecedented level of control of Cas9 activity.

Our observation that the use of Cas9-DD in the absence of Shield-1 leads to the loss of nucleotides on the genomic side of the genome-cassette junction is particularly interesting. One possibility is that these deletions are the result of a transient binding event by Cas9 as it searches for its programmed target site. Cas9 has been found to probe potential binding sites by finding PAMs (Sternberg, Redding *et al*., 2014), and unbiased interrogation of Cas9 cleavage sites in HEK293 cells has revealed that SpCas9 can recognize non-canonical PAMs (Tsai *et al*., 2014b). Additionally, experiments in *C. elegans* have demonstrated that choosing sgRNAs that target loci enriched with potential PAMs increases successful editing (Farboud and Meyer, 2015), which was motivated by the observation that Cas9 can be sequestered by competitor DNA containing a high density of PAM sequences (Sternberg, Redding *et al*., 2014). Those findings, coupled with our observations, suggest that there may be several low-affinity binding events at such target sites that can result in cleavage by Cas9. Of further interest is the co-occurrence of a single C-to-T transition upstream of the PAM with the nucleotide loss event. Because neither allele of JF10 possesses a T at this position and we only observed this mutation when using destabilized Cas9, we speculate that this T could be the result of a deamination of a methylated C, which may occur asymmetrically, as we have only observed this occurrence in the T-C-T allele and not the C-C-C allele. Embryonic stem cells are known to possess non-CpG methylation at CpA and CpT sites, which appears to be mediated in part by Dnmt3a (Ramsahoye *et al*., 2000). Additionally, homologous recombination of direct repeats of GFP facilitated by I-SceI in mouse ESCs has been found to stimulate *de novo* methylation through Dnmt1 that leads to silencing of the recombined cassette (Cuozzo, Porcellini *et al*., 2007). These findings combined with the consistent presence of the mutation when using destabilized Cas9-DD lead us to speculate that DSBs induced by destabilized Cas9-DD could be repaired in a different manner than those of WT-Cas9 and stabilized Cas9-DD. Additionally, the lack of knock-in events when using destabilized DD-Cas9 suggests that DD-Cas9 may be preferable for restricting Cas9 activity. This observation is in accordance with the initial work of Banaszynski *et al*. (2006) describing N-terminal DD fusions as inherently less stable than C-terminal fusions.

Analyzing cells in bulk is informative, but it lacks the resolution only possible at the single-cell (or –clone) level. When working with hiPSCs, single-cell analysis is possible, albeit rather difficult, as the current method for obtaining pure clones is to single-cell sort cells into numerous 96-well plates and analyze the survivors once colonies have formed. The use of a sib-selection-based strategy for enrichment of modified cells is useful, but this strategy is dependent on subsequent subcloning for isolating a pure population as well (Miyaoka *et al*., 2014). Thus, in our analyses, we investigated the consequences of KiBL via single-cell PCR. Cell I was found to have only received the 5’sgRNA vector, but the cassette was knocked-in precisely without loss of any nucleotides, underscoring that KiBL is compatible with single sgRNAs as well. Cell II received both sgRNA vectors, and the modified allele possessed a precise 5’ genomic-imprecise 5’ cassette junction and a 3’ precise cassette-imprecise 3’ genomic junction, which is somewhat consistent with destabilized Cas9-DD targeting a nearby incorrect PAM. Interestingly, in both cells, the other allele was wholly correct except for an additional A at the cleavage site of the 5’ sgRNA. It is possible that this additional A is the result of error-prone NHEJ or that there is a small subpopulation of cells in culture that acquired an A at this position undergoing KiBL. Further analysis involving deep sequencing and more single-cell PCR would have to be carried out to determine which actually occurred. It is worth noting that single-cell cloning is the major obstacle to the application of the CRISPR/Cas9 system to hiPSCs. Additionally, our analysis suggests that multi-sgRNA KiBL may benefit from having both sgRNAs present within the same vector, similar to what has been developed for multiplexed editing using lentiviral vectors (Kabadi *et al*., 2014).

In this work, we chose to identify off-target KiBL events through a targeted, candidate-based method utilizing off-target sites possessing the fewest number of mismatches. While this method provides a good starting point, it does not necessarily take into account the finding that SpCas9 can tolerate bulges in the target DNA and the sgRNA, as well as cleave at non-canonical PAMs and tolerate more than four mismatches (Lin *et al*., 2014; Tsai *et al*., 2014b). Whole genome sequencing would identify the off-target KiBL events, in addition to error-prone off-target cleavage repair, but its cost makes it prohibitive for hiPSCs, and the additional information may not be worth the increased cost, as error-prone off-target repair appears not to occur at high frequency in hiPSCs and primary stem cells (Mandal *et al*., 2014; Yang, Grishin, Wang *et al*., 2014). There are currently three unbiased methods for identifying off-target cleavage events: GUIDE-seq, Digenome-seq, and identification through integrase-deficient lentiviral vectors (Tsai *et al*., 2014b; Kim *et al*., 2015; Wang *et al*., 2015). GUIDE-seq and identification through integrase-deficient lentival vectors are functionally similar, and it does not escape us that KiBL could be used in a similar approach. Ideally, this would be combined with a linear amplification-based method to maximize sensitivity. Such a strategy has been previously used to examine translocations generated by TALENs and CRISPR/Cas9 and to measure the frequency of promoterless gene targeting using adeno-associated virus vectors (Frock *et al*., 2014; Barzel *et al*., 2015).

While being able to place exogenous sequences at any genomic location is useful, being able to knock-in sequences at loci where transgenes do not disrupt endogenous gene regulation (known as safe harbor sites) is also useful. We chose the H11 as our target locus because it has previously been identified as a safe harbor site in the human and mouse genome (Zhu *et al*., 2014; Tasic *et al*., 2011). The use of a safe harbor site is of great importance for the placement of transgenes in cell therapies because of the potential for the integration event itself to perturb endogenous genes in a possibly oncogenic manner (for further information see Sadelain, Papapetrou, and Bushman *et al*., 2011). Additionally, the gene order of the H11 locus appears to be highly conserved, particularly with respect to mammals and through vertebrates in general. For example, the order appears even to be conserved in the coelacanth. This conservation makes H11 particularly attractive as a genome engineering site.

Ultimately, we have presented a method for utilizing the CRISPR/Cas9 system to facilitate *in* vivo blunt-end cloning in precise, homology-independent manner through NHEJ. This method, KiBL, is made possible by the ability of Cas9 to make blunt DSBs, which we and others have exploited using two sgRNAs at once to facilitate precise excision of the intervening sequence (Cong *et al*., 2013; Mali, Yang *et al*., 2013; Canver, Bauer *et al*., 2014; Zheng, Cai *et al*., 2014). The main advantages of our method are its relatively high frequency of precise genomic-cassette junctions, its lack of incorporation of additional exogenous sequences, and its independence from sgRNA efficiency. Additionally, KiBL may prove amenable to high-throughput applications, whereas HR is not, due to the necessity of constructing homology arms for each targeted locus. This construction is of particular concern for the concept of saturation editing, which could become very expensive for targeting large numbers of loci (Findlay, Boyle *et al*., 2014). The main drawbacks of our method are generating large amounts of cassette (which can be easily overcome) and the lower electroporation efficiency of linear dsDNA. The limitation of the amount of cassette generated greatly affects directly comparing KiBL to CRISPR/Cas9-mediated HR, because HR uses at least 10-fold more donor vector than our method (≥ 2μg versus 200ng; Byrne *et al*., 2014). Thus, there is more donor vector available for incorporation in HR relative to KiBL. We recommend that the cassettes be made with unphosphorylated primers each possessing at least three phosphorothioate bonds in the 5’ termini. This property is exemplified in comparing KiBL to the homology-independent knock-in method used in zebrafish (Auer *et al*., 2014). Where we have observed high levels of precise genome-cassette junctions in our hiPSCs, Auer and colleagues (2014) observed only 17% of targeted embryos possessed perfect repair. This decreased level of precise repair may be due to unprotected free ends of the cleaved plasmid, as well as the capability of a perfectly repaired site to be recleaved by the same sgRNA:Cas9 complex. Interestingly, our observation of a lack of 0/0 precise repair when targeting the MYOD1 3’UTR with our PAMs 3’ sgRNA pair (which regenerates a perfect target site for one of the sgRNAs) lends additional support to recleaving by Cas9. This finding, combined with the rest of our analysis of paired sgRNA-mediated deletions and observations by others (Cong *et al*., 2013; Mali, Yang, *et al*., 2013; Canver, Bauer *et al*., 2014; Byrne *et* al., 2014; Ghezraoui *et al*., 2014) also underscores that NHEJ resulting from blunt, chemically unmodified DSBs, such as those made by Cas9, are repaired with a high degree of precision, and that this may be the predominant NHEJ repair mechanism (reviewed in Bétermier, Bertrand, and Lopez, 2014). This observation suggests that the error-prone NHEJ previously observed when only one sgRNA is used may be the result of an alternative repair pathway activated by consistent recleavage of the target site by Cas9. While we have mainly focused on using paired sgRNAs for KiBL, our own data also demonstrate that KiBL can be performed with one sgRNA as well.

KiBL offers a viable alternative to HR and may be a better choice for targeting some cell types. For example, aged hematopoietic stem cells and aged skeletal muscle stem cells retain the capacity to precisely repair DSBs through NHEJ, making KiBL the method of choice for genome engineering in them, particularly *in vivo* or in their quiescent, i.e., non-dividing, state (Flach *et al*., 2014; Beerman *et al*., 2014; Vahidi Ferdousi *et al*., 2014). In summary, KiBL is a versatile method capable of facilitating advanced genome engineering strategies and providing new insights into how Cas9-induced DSBs are repaired.

## Materials and Methods

### Choice of sgRNAs and Vector Construction

All sgRNAs were chosen using the MIT CRISPR Design tool (http://crispr.mit.edu/). Briefly, genomic regions consisting of up to 250 bp were chosen for each locus, and the highest quality guides were chosen for cloning into the pX330 (Addgene #42230) backbone following the protocol developed by Feng Zhang’s lab (https://www.addgene.org/static/cms/filer_public/95/12/951238bb-870a-42da-b2e8-5e38b37d4fe1/zhang_lab_grna_cloning_protocol.pdf). These regions appear in Table 1. All oligos, primers, and gBlocks were ordered from Integrated DNA Technologies (IDT, Coralville, IA). A description of the sgRNAs chosen, as well as the oligos used for cloning, appear in Table 2. sgRNA/Cas9 vectors, as well as all additional plasmids in this study, were isolated using the Nucleobond Midi Plus EF kit (Machrey-Nagel, Bethlehem, PA).

**Table 1.** Interrogated regions used for design of sgRNAs. The GRCh37/hg19 assembly was used for human regions and the GRC38/mm10 assembly was used for murine regions.

| Gene | Region # | Chromosomal Location |
| --- | --- | --- |
| *MYOD1* | 1 | Hs chr11: 17742664-913 |
| *MYOD1* | 2 | Hs chr11: 17743679-928 |
| *PAX3* | 1 | Hs chr2: 223066487-736 |
| *PAX7* | 1 | Hs chr1: 19062359-608 |
| *PAX7* | 2 | Hs chr1: 19062633-845 |
| *PAX7* | 3 | Hs chr1: 19061876-2125 |
| *mUtrn* | 2 | Mm chr10: 12381712-961 |
| *mUtrn* | 4 | Mm chr10: 12384485-734 |
| H11 locus | 1 | Hs chr22: 31830203-452 |
| H11 locus | 2 | Hs chr22: 31830847-1096 |

**Table 2.**
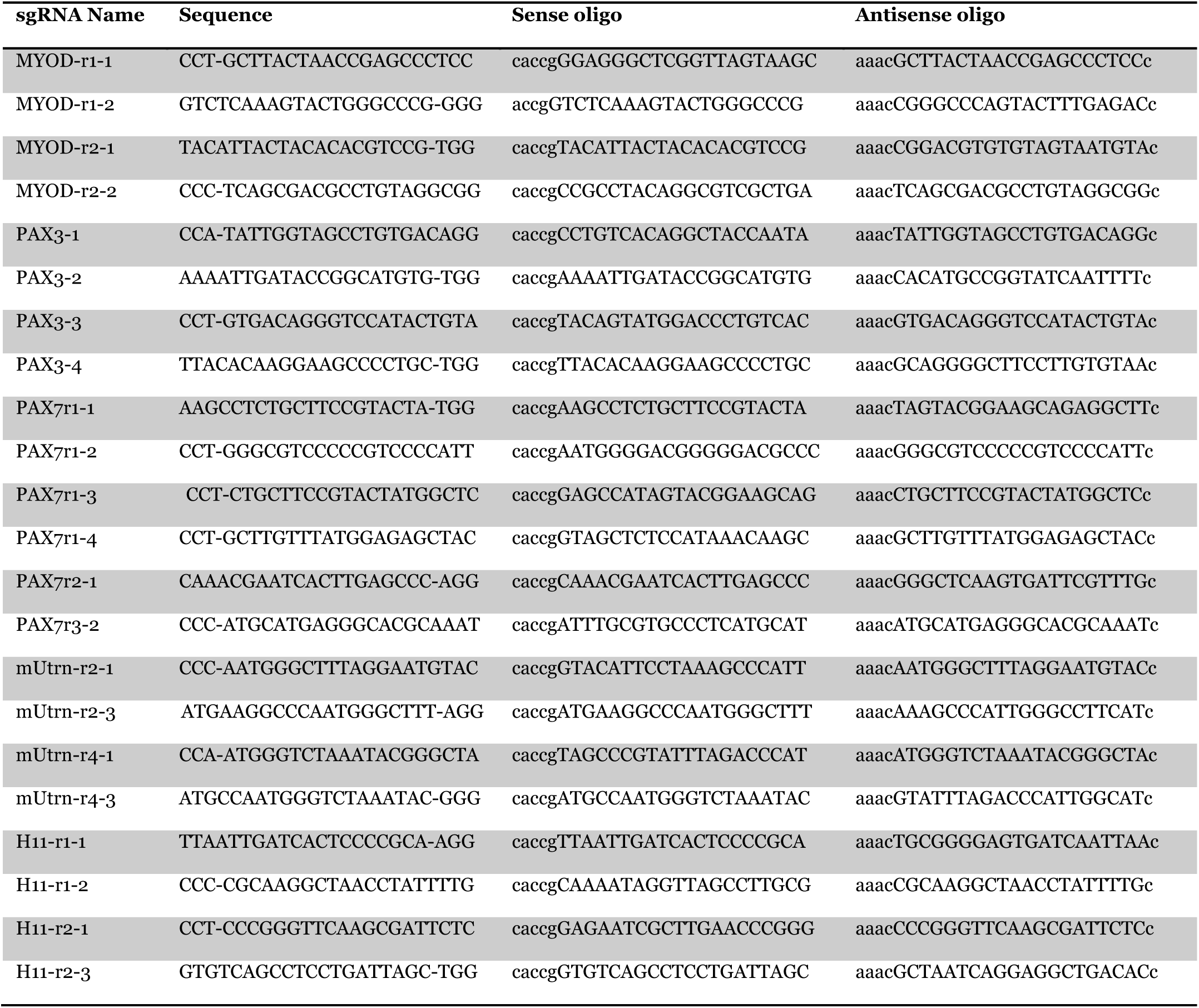
sgRNAs and cloning oligos used. All sequences are in the 5’ to 3’ direction and relative to the sense strand where appropriate. Dashes separate PAM from target. Lower case indicates overhang sequence for cloning.

The C-terminal L106P destabilization domain was obtained as a gBlock (IDT), and was digested with *Fse*I (New England Biolabs (NEB), Ipswich, MA) and *Bsa*I-HF (NEB). pX330 was digested with *Fse*I and *Eco*RI-HF (NEB), and dephosphorylated with Antarctic phosphatase (NEB). The digested C-terminal DD and the pX330 backbone were ligated with T4 DNA ligase (NEB), and transformed into STBL3 chemicompetent cells (Life Technologies, Grand Island, NY). In order to clone sgRNAs into pX330-Cas9-DD, the *Bbs*I cloning site was replaced with a *Bsa*I x2 cloning site using a gBlock containing the entire sgRNA expression cassette. Both pX330-Cas9-DD and the gBlock were sequentially digested with *Pci*I (NEB) and *Kpn*I-HF (NEB) before dephosphorylation of the backbone and subsequent ligation and transformation to generate pX330-*Bsa*Ix2-Cas9-DD. This new cloning site uses FastDigest Eco31I (isochizomer of BsaI; Thermo Fisher Scientific, Waltham, MA) and the two following general primers:

Sense oligo: 5’-accg-NNNNNNNNNNNNNNNNNNNN-3’ Antisense oligo: 5’-aaac-NNNNNNNNNNNNNNNNNNNN-3’

Additionally, pX330-Cas9-DD versions of the H11 sgRNAs were generated by cloning Cas9-DD directly into their pX330 vectors through the use of FastDigest *Bsh*TI (Thermo Fisher Scientific) and FastDigest *Not*I (Thermo Fisher Scientific).

To generate pX330-BsaIx2-DD-Cas9, the N-terminal DD was obtained as a gBlock and digested with FastDigest *Bsh*TI and FastDigest *Bgl*II (Thermo Fisher Scientific) before being cloned into the pX330 backbone that had been digested with the same enzymes and dephosphorylated with FastAP (Thermo Fisher Scientific). This vector was subsequently digested with FastDigest *Kpn*I (Thermo Fisher Scientific) and FastDigest *Not*I to isolate the DD-Cas9 insert. pX330-*Bsa*Ix2-Cas9-DD was digested with the same enzymes and dephosphorylated to remove Cas9-DD. The DD-Cas9 insert was cloned into the pX330-BsaIx2 backbone to generate pX330-BsaIx2-DD-Cas9.

### Knock-In Cassette Construction

The DICE-EPv2.0 and PTKmChR cassettes were amplified from a common vector containing the Puro^R^ΔTK fusion gene (from Addgene # 22733) followed by a P2A ribosomal skipping element and mCherry followed by the rabbit beta-globin terminator. This cassette is under the control of the mouse phosphoglycerol kinase (PGK) promoter. Cassettes were amplified via PCR with Q5 high-fidelity polymerase, subjected to gel electrophoresis, excised, and purified with the MinElute Gel Extraction Kit. For the DICE-EPv2.0 cassettes, the phiC31 and Bxb1 *attP* sites were included in the primers used for amplification. All primers used for cassette amplification appear in Table 3.

**Table 3.**
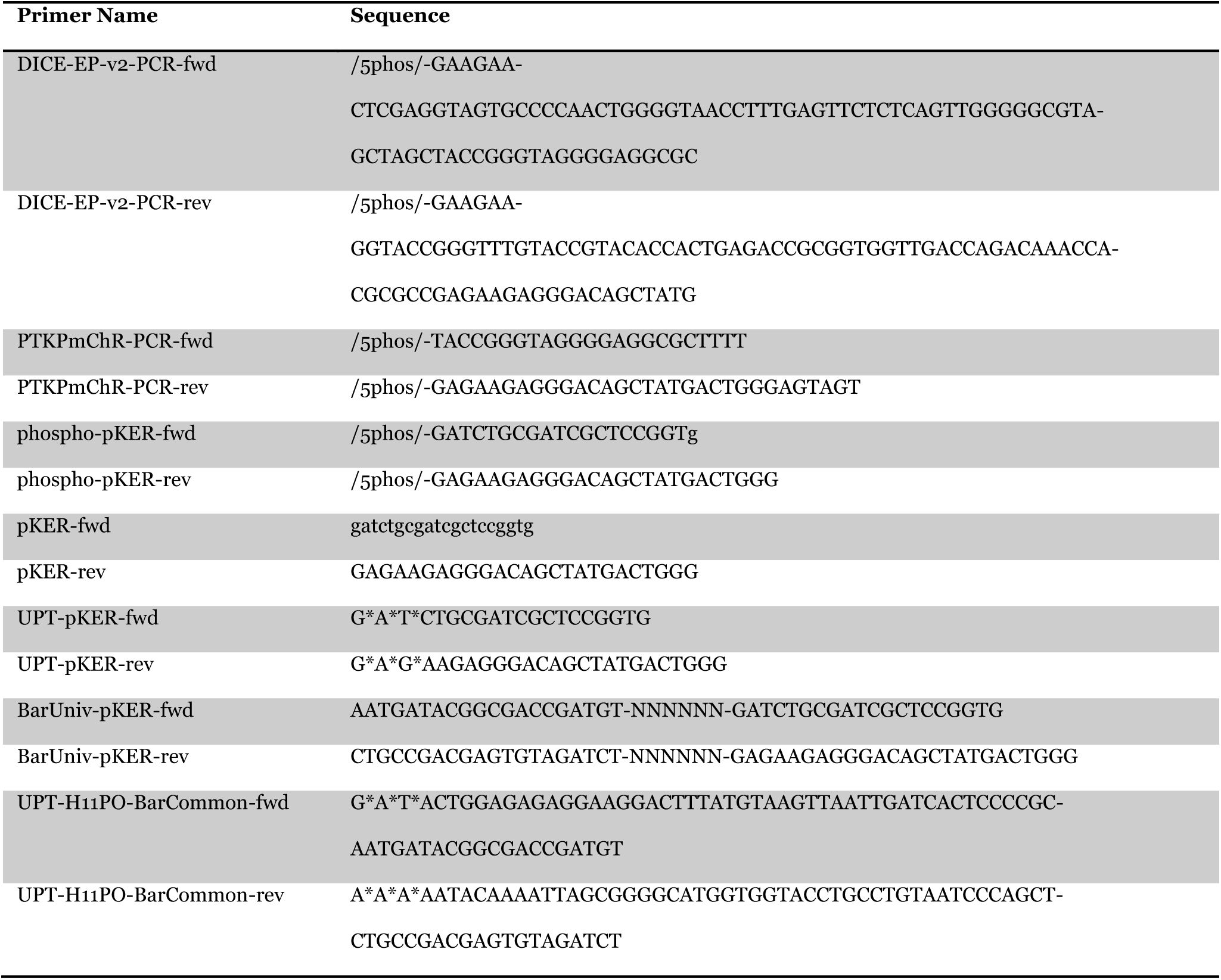
Primers used in cassette amplification. All oligos are in the 5’ to 3’ direction.

The pKER cassettes were assembled by first obtaining phosphorylated gBlocks for Clover, mRuby2, and mCerulean in the form of EF1α promoter-fluorescent protein-rabbit beta globin terminator. These three gBlocks were cloned separately into a kanamycin resistance backbone with blunt ends. mCardinal and mKOκ were obtained as gBlocks, digested with FastDigest NheI (Thermo Fisher Scientific) and FastDigest NotI, and cloned into a dephoshorylated pKER-Clover backbone that had been digested with the same enzymes to remove Clover. Cassettes were amplified from 1-3 ng of plasmid using Q5 high fidelity polymerase in eight individual reactions. After amplification, the reactions were pooled and purified using either the MinElute PCR Purification kit (Qiagen) or the GeneJet PCR Purification kit (Thermo Fisher Scientific). The purified reaction was then treated with FastDigest *Dpn*I to digest residual plasmid. After digestion, the reaction was brought to 100 μL with nuclease-free water (Life Technologies) and purified with a CHROMASpin-1000+TE chromatography column (Clontech, Mountain View, CA) to remove digested plasmid and restriction enzyme. Barcoded cassettes containing homology arms were first amplified with primers containing, in the 5’direction, a 20-base secondary priming sequence, a 6-base randomized barcode, and the priming sequence for the pKER cassette. This reaction was then purified with the GeneJet PCR Purification kit and used as the template for secondary PCR using primers containing 50-base homology arms and the secondary priming sequence, which was purified and digested in the same manner as described above. All primers used for cassette amplification appear in Table 3.

### Cell Culture

HEK293 cells were maintained in 6-well tissue culture plates coated with Poly-L-Lysine (Sigma-Aldrich, St. Louis, MO) in high-glucose DMEM (Thermo Fisher Scientific) supplemented with 10% fetal bovine serum (FBS; Gemini Bio-Products, West Sacramento, CA), 1x non-essential amino acids (Life Technologies), and 1x GlutaMAX (Life Technologies). Cells were passaged with 0.05% Trypsin-EDTA (Life Technologies) as needed at a 1:12 split. C2C12 cells were maintained in the same media as HEK293 cells and passaged in the same manner. C2C12 cells were grown on 6-well tissue culture plates. H9 hESCs were maintained in 0.1% gelatin-coated 6-well tissue culture plates on β-irradiated mouse embryonic fibroblast feeder cells in human embryonic stem cell media consisting of DMEM/F12 (Thermo Fisher Scientific), 20% Knockout Serum Replacement (Life Technologies), 1x non-essential amino acids (Life Technologies), 1x GlutaMAX (Life Technologies), 0.1mM β-mercaptoethanol (Life Technologies), and 8ng/mL bFGF (PeproTech, Rocky Hill, NJ). Cells were passaged as clumps using collagenase IV (Stem Cell Technologies, Vancouver, British Columbia, Canada). JF10 hiPSCs were maintained in 6-well tissue culture plates on recombinant human vitronectin (Life Technologies) in Essential 8 media (Life Technologies). Cells were passaged as clumps every three to four days with Versene (Life Technologies) at a 1:6 split.

### Detection of CRISPR/Cas9-Facilitated Excision

HEK293 and C2C12 cells were transfected using FuGene HD (Promega, Madison, WI). Briefly, 150,000 cells were plated one to two days prior on poly-L-lysine-coated (HEK293) or standard (C2C12) 24-well tissue culture plates and were transfected with 5 μg (HEK293) or 3 μg (C2C12) of each sgRNA/Cas9 vector for a total of 10 μg or 6 μg per transfection, respectively, in Opti-MEM I (Life Technologies) media with FuGene HD being used at a 3:1 ratio. Cells were harvested four days post-tranfection and genomic DNA was isolated using the DNeasy Blood and Tissue Mini Kit (Qiagen). Each transfection was separately performed at least twice. H9 hESCs were transfected using the Amaxa Nucleofector (Lonza, Basel, Switzerland) with the Human Embryonic Stem Cell Nucleofection Kit 2 (Lonza). Briefly, cells were passaged normally into Matrigel-coated 6-well plates three days prior to nucleofection. Before nucleofection, media was replaced at least 1 hr beforehand with fresh media containing 10μM ROCK inhibitor Y-27632 (R&D Systems, Minneapolis, MN). Cells were dissociated with collagenase IV, and subsequently electroporated with 1.5 μg of each sgRNA/vector for a total of 3 μg using program B-16 according to the manufacturer’s instructions, and plated on 0.1% gelatin-coated 12-well plates containing at one reaction per well in human stem cell media containing 10μM ROCK inhibitor Y-27632. Each reaction was carried out at least twice Genomic DNA was isolated from the cells 2 days later with the DNeasy Blood and Tissue Mini Kit.

PCR amplification of the deletion-containing regions was carried out using OneTaq (NEB) and Q5 high fidelity polymerase using 150 ng of gDNA according to the manufacturer’s instructions. Amplicons were subjected agarose gel electrophoresis, and subsequently the deletion-containing bands, as well as the full-length bands where applicable, were excised and purified using the MinElute Gel Extraction Kit (Qiagen). Purified amplicons were then subcloned using the CloneJet PCR Cloning Kit (Thermo-Fisher Scientific), and subsequently transformed into α-Select electrocompetent cells (Bioline, Taunton, MA), 10-β chemicompetent cells (NEB), or STBL3 chemicompetent cells according to their manufacturers’ instructions. Colonies were inoculated and grown overnight. Plasmids were isolated the following day using the Miniprep Spin Kit (Qiagen), and subsequently subjected to Sanger sequencing of the inserts. Primers used for amplification of excision events appear in Table 4.

**Table 4.**
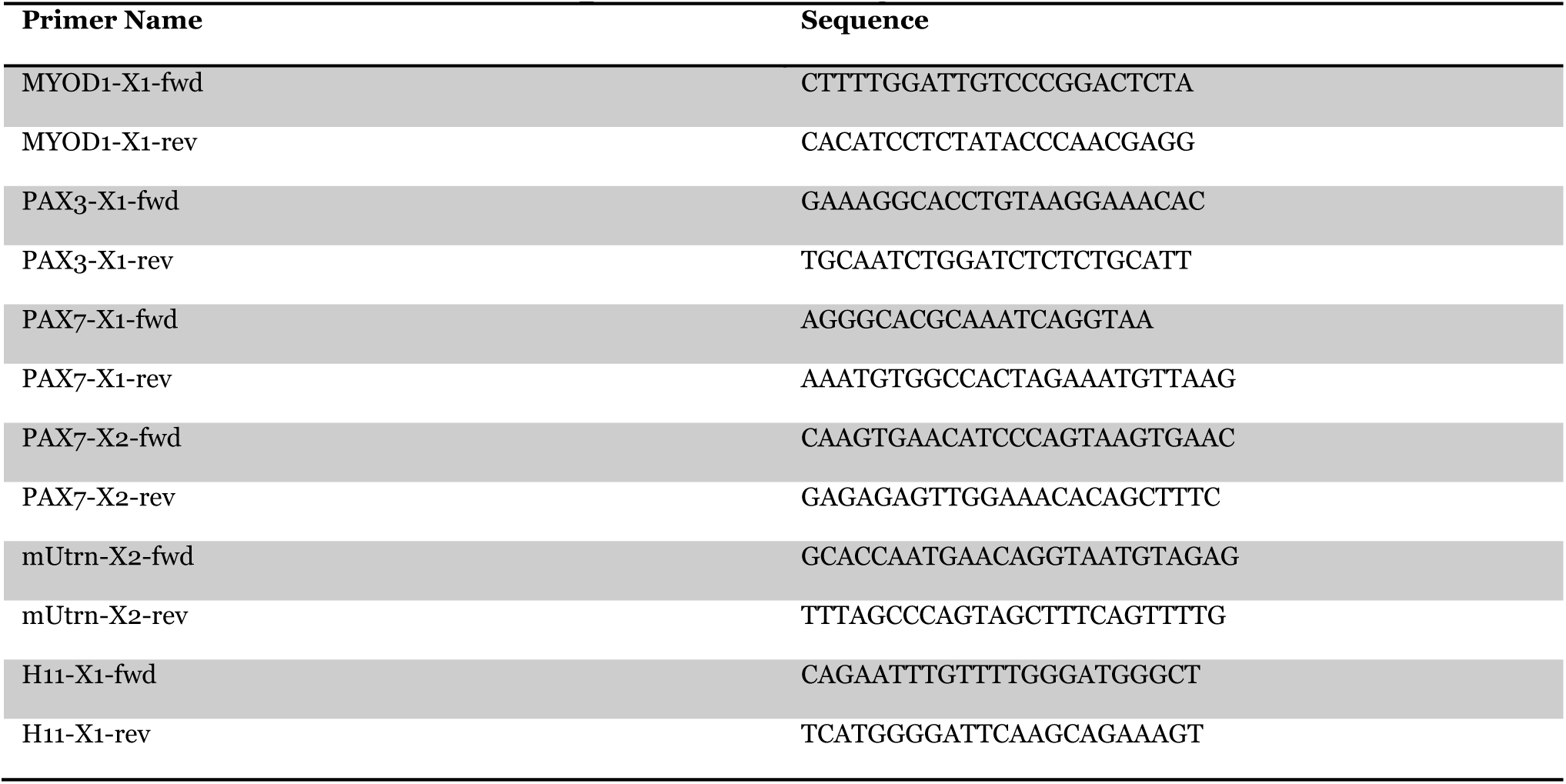
Excision detection primers. All oligos are in the 5’ to 3’ direction.

### Flow Cytometry

Flow cytometric analysis was carried out on LSRII- and FACScan-class analyzers (BD Biosciences, San Jose, CA). Sorting was carried out on FACSAriaII-class sorters (BD Biosciences). Live cells were discriminated on the basis of DAPI exclusion using the NucBlue Fixed Cell Stain ReadyProbes reagent (Life Technologies). mCherry was detected via excitement with a 561 nm yellow/green 100 mW laser with a 571 nm LP filter in the optical path, a 725 nm SP splitter, and a 690±40 nm BP filter. Clover was detected via excitement with a 488 nm 50mW blue laser, a 505 nm LP splitter, and 525±50 nm BP filter. mKOκ was detected via excitement with a 532 nm 150mW green laser and a 575±25 nm BP filter. mCardinal was detected with a 640 nm 40mW red laser and a 670±30 nm BP filter. mRuby2 was detected with a 532 nm 150mW green laser, a 600 nm LP splitter, and a 610±20 nm BP filter. mCerulean was detected with a 405 nm 50mW violet laser and a 450±50 nm BP filter in the absence of DAPI. All flow cytometric data were analyzed using FlowJo software (Tree Star, Ashland, OR).

### Detection of Knock-in Blunt Ligation and Homologous Recombination

For HEK293 cells, FuGene HD was used to transfect cells as described above, using 100 ng of cassette and 1.5 μg of each sgRNA/Cas9 vector for a total of 3.1 μg of DNA. 500,000 cells had been plated one to two days prior to transfection. Reactions were carried out in triplicate. Shield-1 (Clontech) was added to the cells immediately prior to transfection at a concentration of 1 μM. Puromycin (Life Technologies) selection was started two days after tranfection at a concentration of 1μg/mL and carried out for four days with fresh media and antibiotic being replaced every two days. Cells were analyzed by flow cytometry two days after transfection or six days in the case of puromycin-selected cells. Briefly, cells were trypsinzed with 0.05% trypsin, resuspended in PBS (Thermo Fisher Scientific) with 2% FBS, and were filtered before analysis to remove clumps and debris. gDNA was isolated from the remaining cells with the DNeasy Blood and Tissue Mini Kit. Genome-cassette junctions were amplified with Q5 Hi-Fidelity polymerase using at least 100 ng of DNA per reaction. Amplicons were subjected to gel electrophoresis, excised, purified, subcloned, transformed, and sequenced as described above. Primers used appear in Table 5.

**Table 5.**
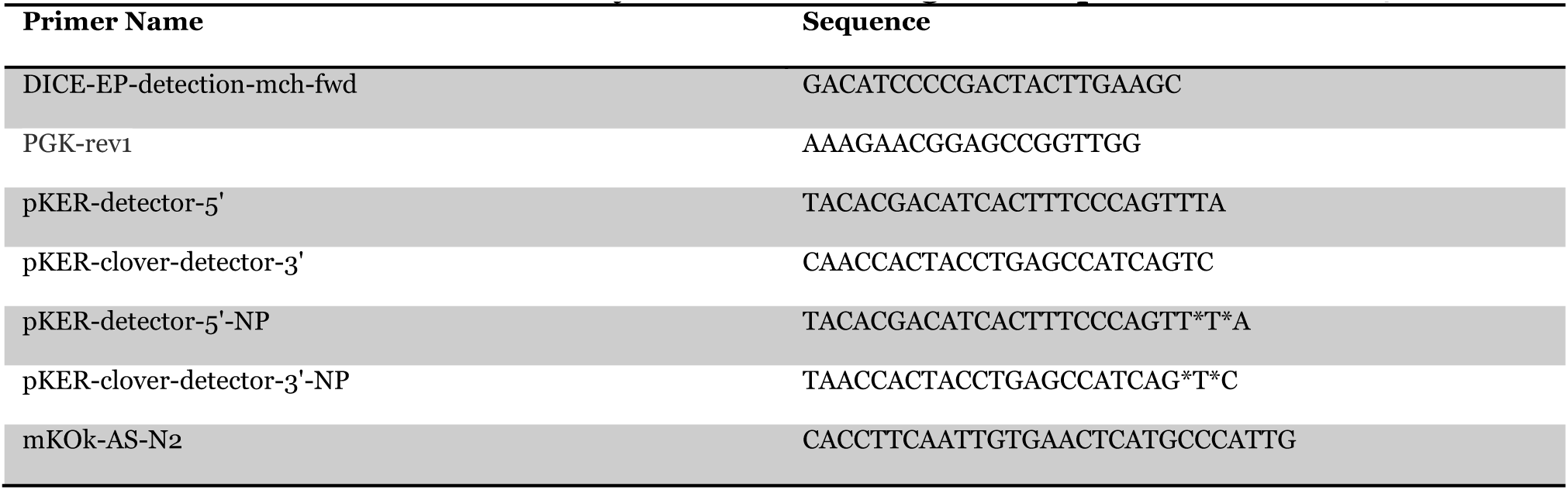
Primers for Amplifying Cassette Junctions. All oligos are in the 5’ to 3’ direction and are to be used in conjunction with the genomic primers in Table 4.

JF10 cells were plated three to four days prior to electroporation on vitronectin-coated 6-well plates. Cells were incubated at least one hour beforehand with Essential 8 media containing 10 μM ROCK inhibitor Y-27632. Cells were then dissociated into single cells with Versene. Cells were resuspended in the homemade nucleofector buffer described in Zhang, Vanoli *et al*., 2014, with 500ng of each sgRNA/Cas9 vector and 200ng of cassette per reaction (for a total of 1.2 μg of DNA) and electroporated with the Amaxa nucleofector using program B-16 in triplicate. Electroporated cells were plated on vitronectin-coated 12-well plates in Essential 8 media containing 10 μM ROCK inhibitor Y-27632 and 0, 0.5, or 1 μM of Shield-1. ROCK inhibitor was added each day for four days after nucleofection. Modified cells were subjected to FACS four to seven days after nucleofection. Briefly, cells were treated 1 hour prior with 10 μM ROCK inhibitor Y-27632 before being trypsinized with TrypLE (Life Technologies) for 10 minutes. Harvested cells were resuspended in DPBS (Life Technologies) with 2% AlbuMAX I (Life Technologies), 2 mM EDTA (Life Technologies), NucBlue Fixed Cell Stain ReadyProbes reagent, and 10 μM ROCK inhibitor Y-27632, and subsequently filtered to remove clumps and debris. Genomic DNA was isolated from sorted cells using the ZymoBead Genomic DNA kit, (Zymo Research, Irvine, CA). gDNA was then subjected to targeted amplification of the H11 locus via PCR amplification with Q5 high-fidelity polymerase or multiple displacement amplification (MDA; a variant of whole genome amplification) following Dean *et al*., 2002, with the addition of inorganic (yeast) pyrophosphatase (NEB). PCR-amplified products were purified with the MinElute PCR Purification kit (Qiagen), diluted to 100 μL with Buffer EB (Qiagen), and used for further genome-cassette junction PCR amplifications. MDA reactions were diluted to 200 μL with TE-EF buffer (Machrey-Nagel) and also used for further genome-cassette junction PCR amplifications. Primers used appear in Table 5.

### Analysis of Off-Target KiBL Events

For each sgRNA, forward and reverse primers were designed to each of the top six off-target sites as predicted by the MIT CRISPR Design Tool. Each forward and reverse primer was used in conjunction with the 5’ and 3’ pKER-Clover detection primers, for a total of 48 reactions per sample. 1 μL of 1:200 diluted WGA of Clover+ cells was used in each PCR amplification. PCR was carried out with Q5 high-fidelity polymerase. Off-target primers appear in Table 6.

**Table 6.**
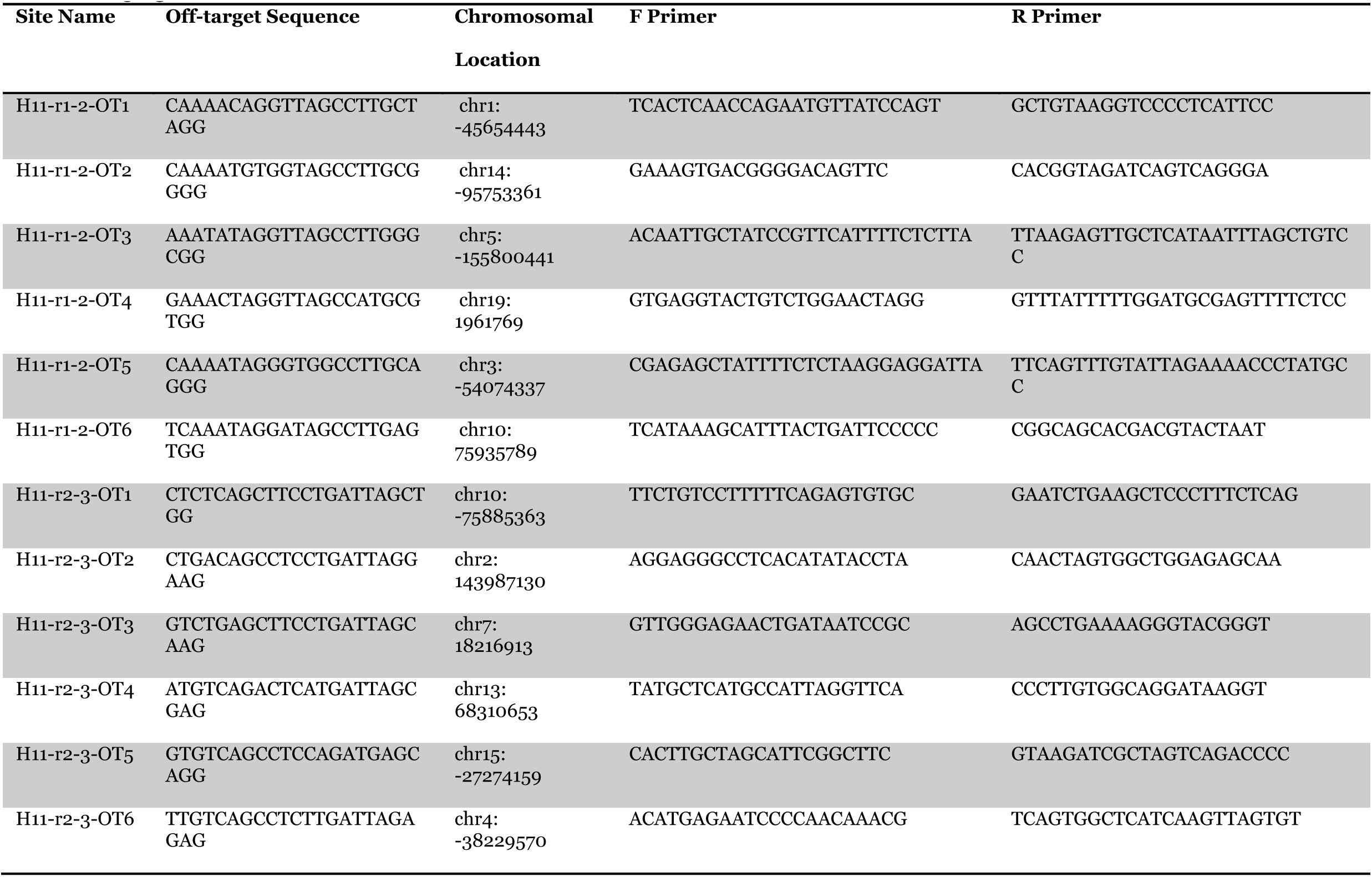
Off-target sites and primers for H11-r1-2 and H11-r2-3. All oligos are in the 5’-3’ direction.

### Western Blotting

HEK293 cells in 6-well plates that had been plated two days prior were transfected using FuGene HD with 5 μg of plasmid encoding the H11-r1-2 sgRNA and WT-Cas9, Cas9-DD, or DD-Cas9 as described above. Cas9-DD and DD-Cas9 were transfected in duplicate. All wells received 0.5 μM Shield-1 except one replicate each for Cas9-DD and DD-Cas9. The following day, media was aspirated and replaced with fresh media containing 0.5 μM Shield-1 for all wells except for the replicates that did not receive Shield-1 the previous day. 5 hours later, media was aspirated, cells were washed twice with cold PBS, and protein was extracted with RIPA buffer (Thermo Fisher Scientific) containing 1x HALT protease inhibitor cocktail (Thermo Fisher Scientific). Cell lysate was snap-frozen for later analysis. Protein concentration was determined with the Bradford assay.

For Western blotting, 50μg of protein was denatured in the presence of 1x Laemmli buffer (Bio-Rad Laboratories, Hercules, CA) and 0.1 M DTT (Life Technologies) at 100°C for 5 minutes. Samples were then resolved on a 10% SDS-PAGE gel (Bio-Rad) at 100 V in 1x Tris/Glycine/SDS buffer (Bio-Rad). Protein was transferred to an Immuno-Blot PVDF membrane (Bio-Rad) under wet transfer conditions at 200 mA for 1 hour with constant current at 4°C in 1x Tris/Glycine/SDS buffer + 10% methanol. Following transfer, the membrane was blocked for 1 hour at room temperature with agitation with Protein-Free T20 blocking buffer (Thermo Fisher Scientific) and incubated overnight with agitation at 4°C with 1:10,000 monoclonal ANTI-FLAG M2 antibody (Sigma-Aldrich) and 1:10,000 monoclonal anti-GAPDH antibody (clone GAPDH-71.1; Sigma-Aldrich). The following day, the membrane was washed three times for 5 minutes each with PBS + 0.05% TWEEN-20 (Sigma-Aldrich). Secondary incubation was carried out at room temperature with agitation for 1 hour with 1:5000 goat anti-mouse IgG conjugated to horseradish peroxidase (Thermo Fisher Scientific). The membrane was washed again as described above and then incubated for 5 minutes at room temperature with Clarity Western ECL substrate (Bio-Rad). Amersham Hyperfilm ECL (GE Healthcare, Buckhamshire, UK) was exposed to the membrane and subsequently developed. The blot was then quantified using ImageJ and normalized to the WT-Cas9 sample.

### Statistics

All statistical analysis was carried out using GraphPad Prism 6.0 (GraphPad Software, La Jolla, CA)

## Acknowledgments

We thank Jennifer Sullivan for designing the off-target primers. We thank Chris Bjornson for comments on and critical reading of the manuscript. We thank Michael Wilkinson for comments and discussion. We also thank the Jain Foundation and Cellular Dynamics Inc. for providing us with the JF10 hiPSCs. Additionally, we thank the staff of the Stanford Shared FACS Facility, particularly Cathy Crumpton and Ometa Herman for sharing their expertise.

---

**Figure 1-figure supplement 1.**
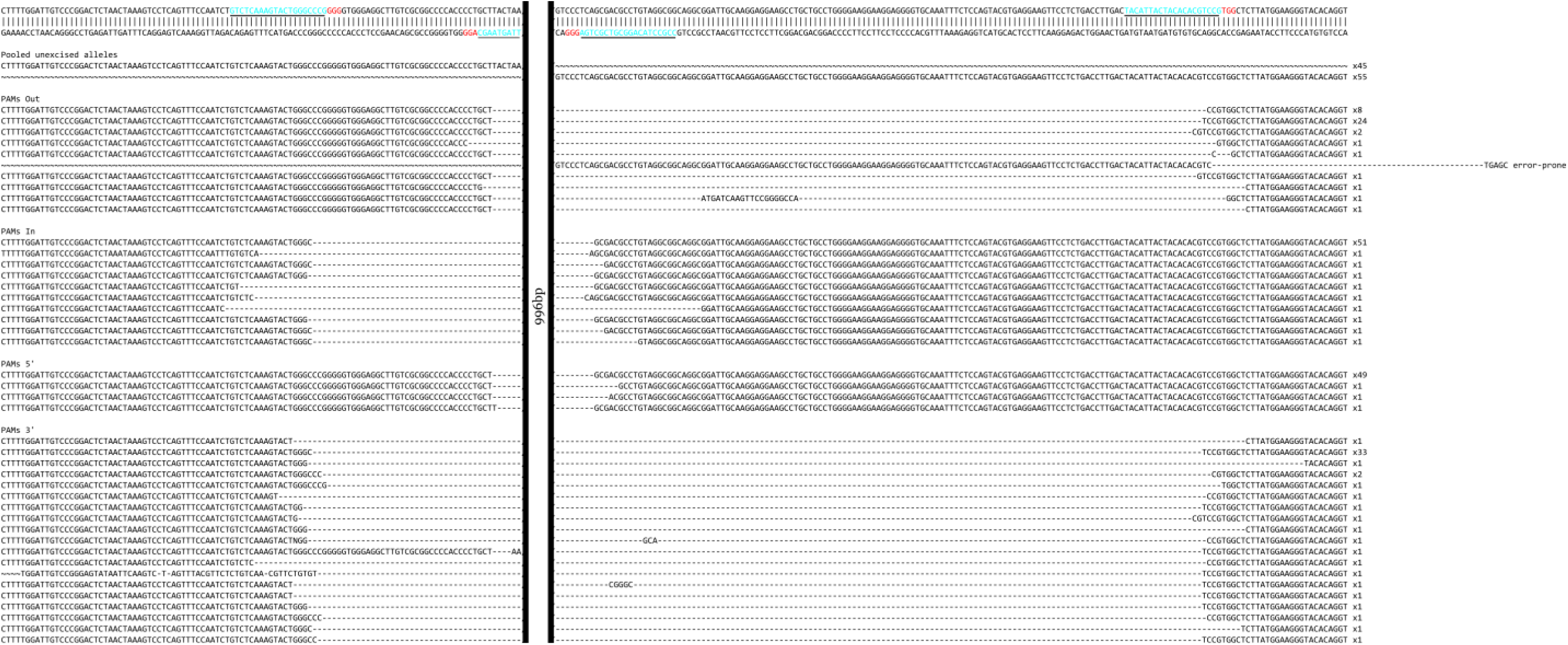
Sequences of CRISPR/Cas9-mediated deletions at the MYOD1 3’UTR. Underlined blue sequence denotes sgRNA targets. Red denotes PAMs. ∼ = Not sequences

**Figure 1-figure supplement 2.**
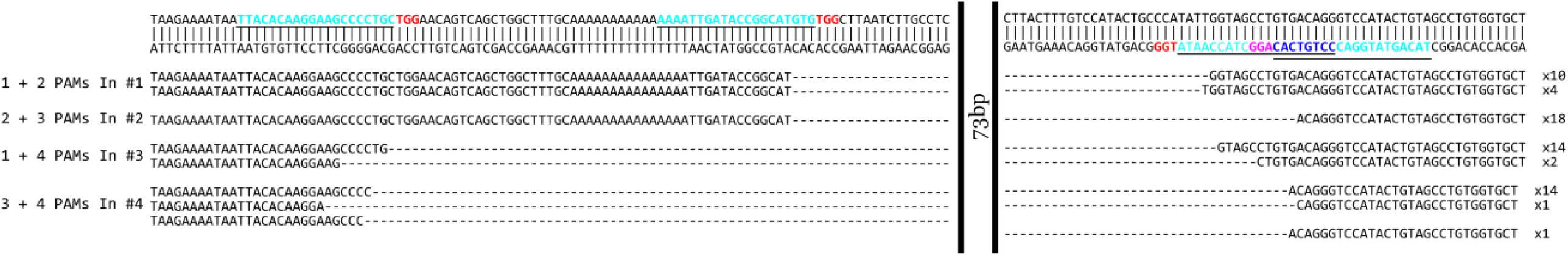
Sequences of CRISPR/Cas9-mediated deletions at the PAX3 3’UTR. Underlined blue sequence denotes sgRNA targets. Red denotes PAMs. Dark blue and pink denote overlapping sgRNA target sequence and PAMs respectively.

**Figure 1-figure supplement 3.**
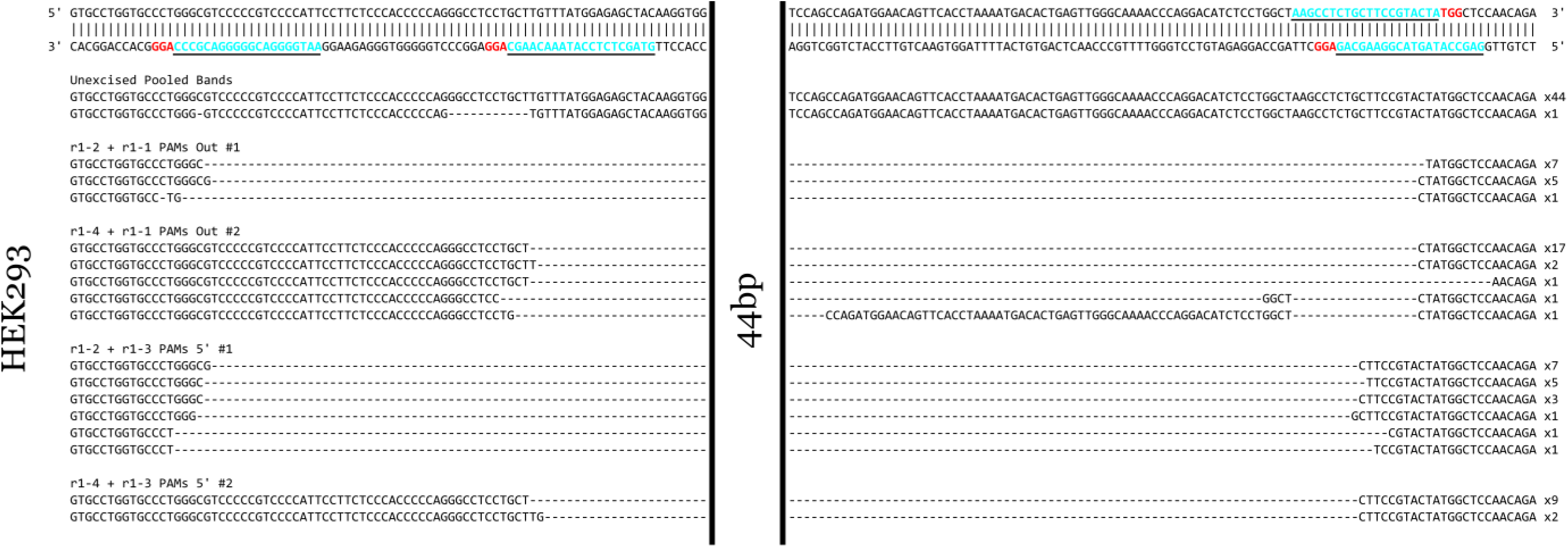
Sequences of CRISPR/Cas9-mediated deletions at the PAX7 3’UTR in HEK293 cells. Underlined blue sequence denotes sgRNA targets. Red denotes PAMs.

**Figure 1-figure supplement 4.**
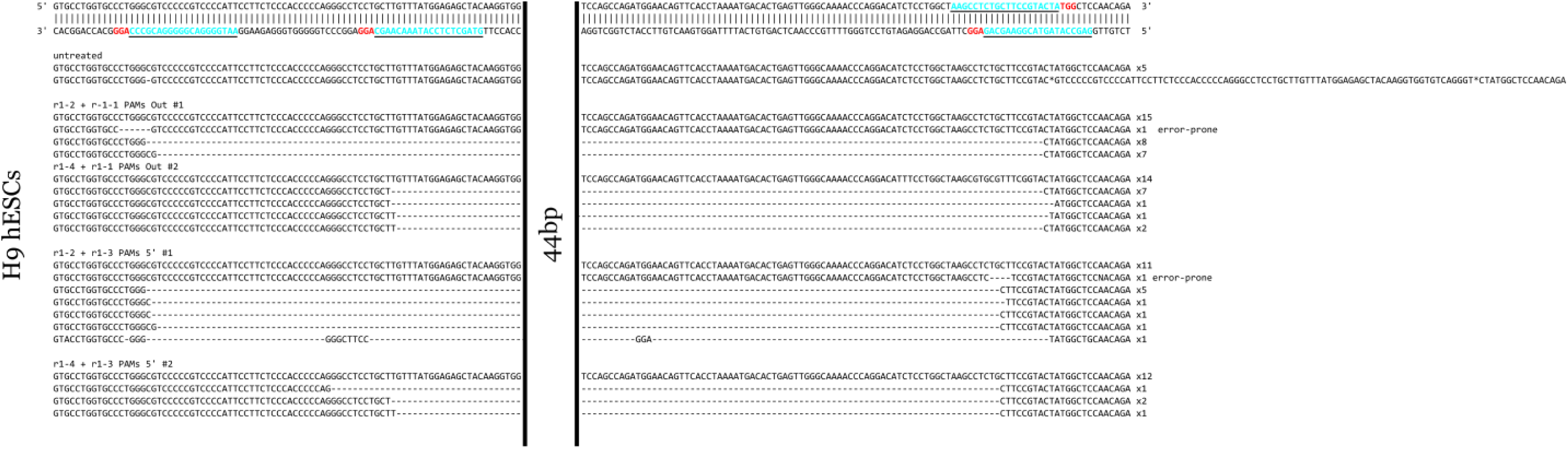
Sequences of CRISPR/Cas9-mediated deletions at the PAX7 3’UTR in H9 hESCs. Underlined blue sequence denotes sgRNA targets. Red denotes PAMs.

**Figure 1-figure supplement 5.**
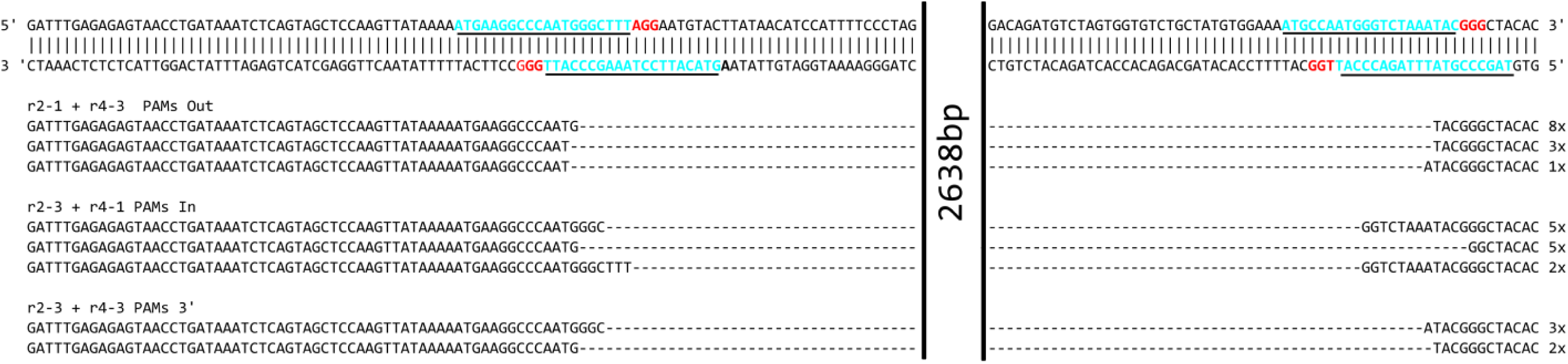
Sequences of CRISPR/Cas9-mediated deletions at the mUtrn 3’UTR. Underlined blue sequence denotes sgRNA targets. Red denotes PAMs.

**Figure 2-figure supplement 1.**
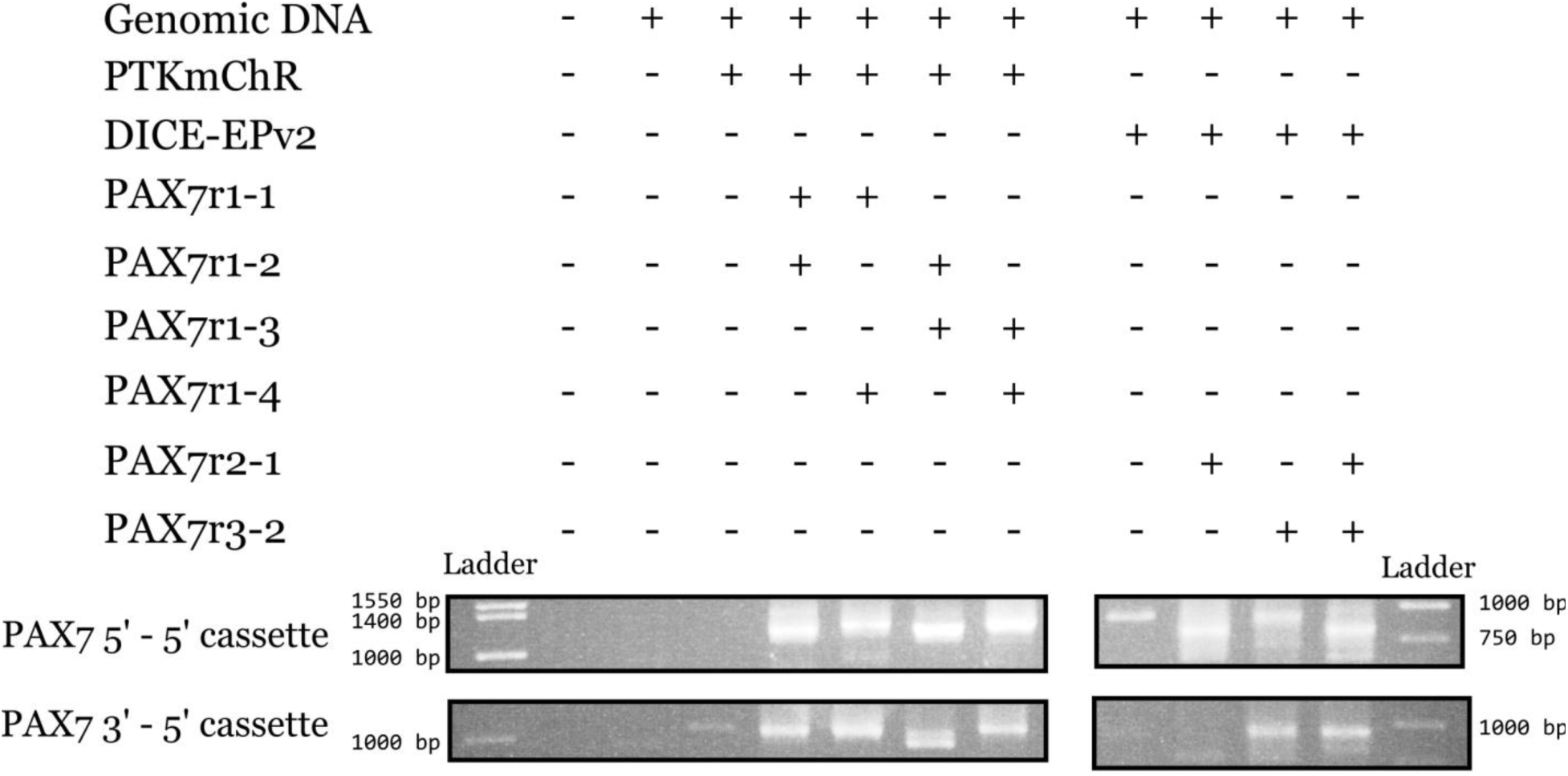
Analysis of PAX7-PTKmChR/DICE-EPv2.0 junctions. PCRs were carried out to analyze the PAX7 5’-5’ cassette and PAX7 3’-5’ cassette junctions in transfected HEK293 cells through agarose gel electrophoresis. The PAX7-X2 primers were used along with the PGK-rev primer for amplification. Marker ladder bands are denoted for reference. Expected band sizes for 5’-5’ junctions: r1-1 + r1-2 = 1188 bp; r1-1 + r1-4 = 1233 bp; r1-2 + r1-3 =1188 bp; r1-3 + r1-4 = 1233 bp; r2-1 = 1595 bp; r3-2 = 808 bp; r2-1 + r3-2 = 808 bp. Expected band sizes for 3’-5’ junctions: r1-1 + r1-2 = 1028 bp; r1-1 + r1-4 = 1028 bp; r1-2 + r1-3 =1036 bp; r1-3 + r1-4 = 1036 bp; r2-1 = 929 bp; r3-2 = 1716 bp; r2-1 + r3-2 = 929 bp.

**Figure 3-figure supplement 1.**
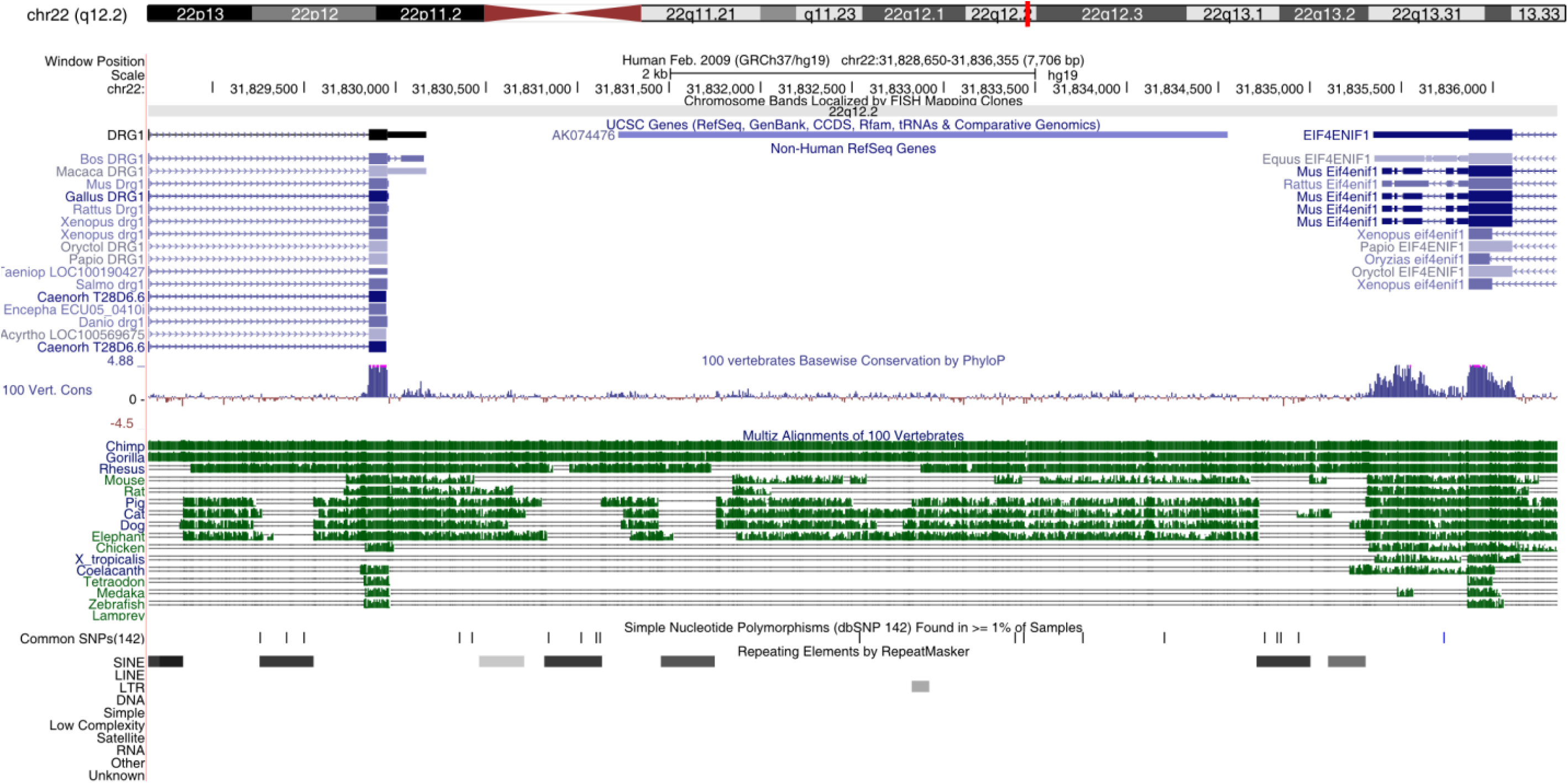
Stucture and conservation of the H11 safe harbor locus. Schematic from the UCSC Genome Browser (http://genome.ucsc.edu; used Feb. 2009 (GRCh37/hg19) assembly) displaying H11 locus in the human genome on chromosome 22. The flanking genes are *DRG1 and EIF4ENIF1*. AK074476 is a potential non-coding RNA isolated from human lung. Below are depicted alignments of the region for several vertebrates. Note the high degree of sequence conservation among mammals and the degree of the gene structure conservation among vertebrates.

**Figure 4-figure supplement 1.**
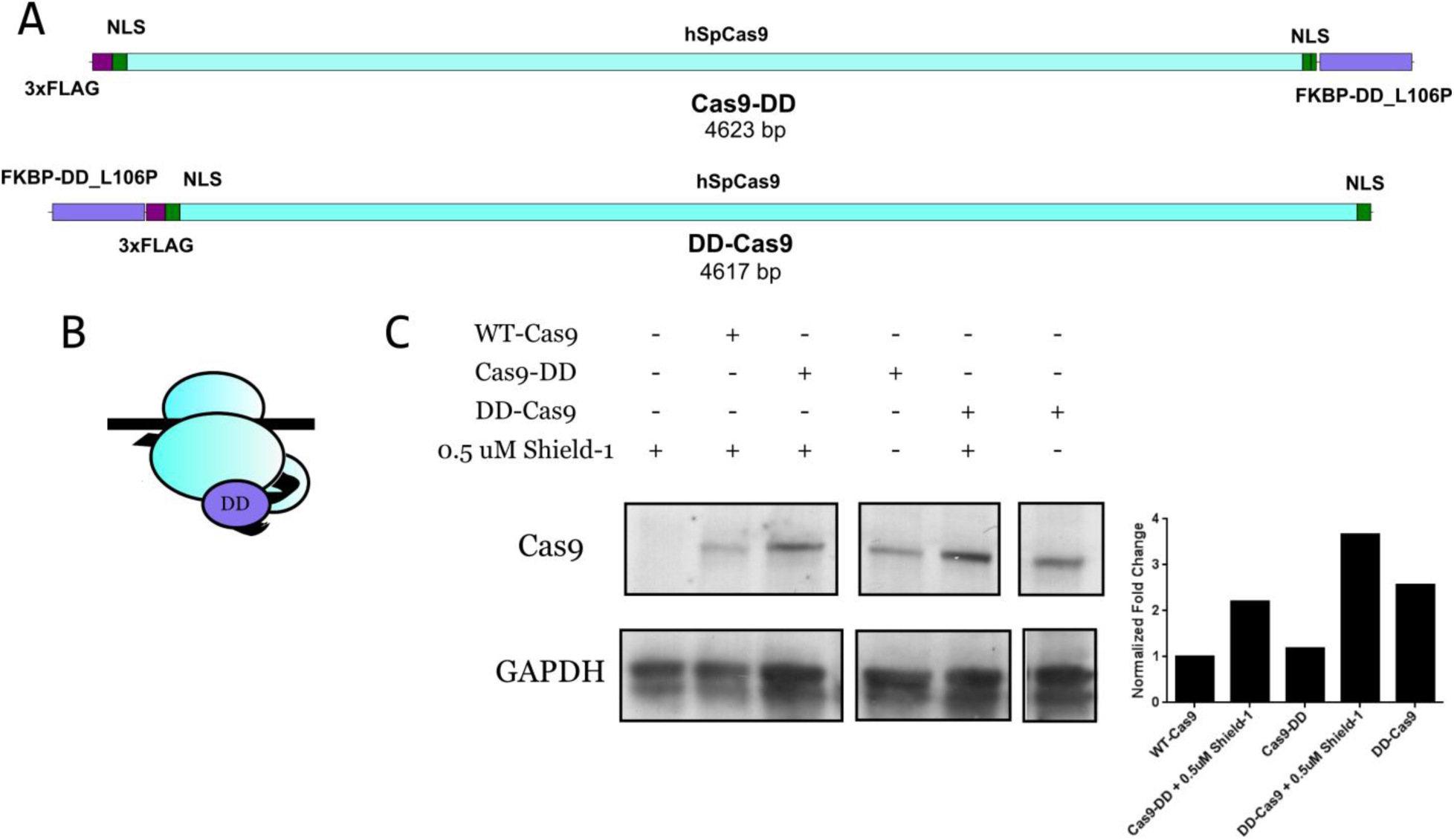
Stability analysis of Cas9-DD and DD-Cas9 in HEK293 cells. (A) Structure of Cas9-DD and DD-Cas9. (B) Schematic of DD-Cas9. (C) Western blot of stabilized and destabilized Cas9-DD and DD-Cas9 in HEK293 cells. Cells were transiently transfected with vectors encoding sgRNA H11-r1-2 and wild-type Cas9, Cas9-DD, or DD-Cas9, in the presence or absence of 0.5 μM Shield-1 for 1 day post transfection. All versions of Cas9 were detected with an anti-Flag antibody. GAPDH serves as a loading control. Quantification of protein levels as relative fold change normalized to wild-type Cas9 appears to the right.

**Figure 4-figure supplement 2.**
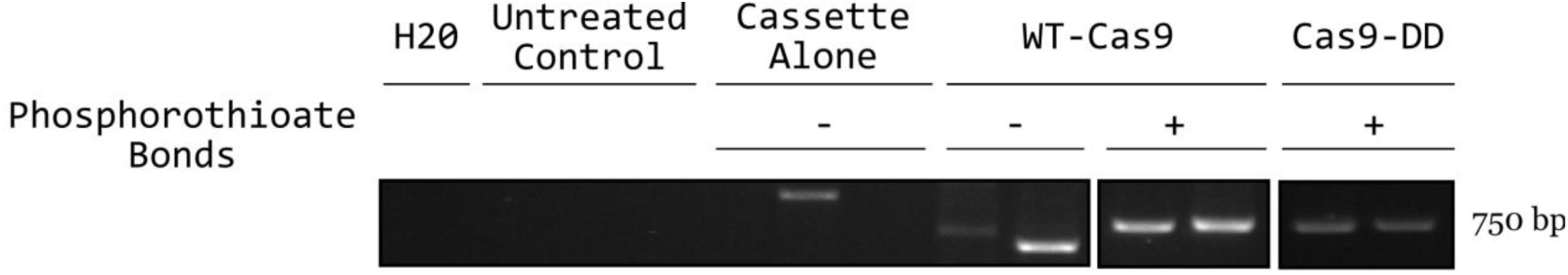
Analysis of the effect of phophorothioate bonds on KiBL. PCR was carried out to analyze the PAX7 5’-5’ pKER cassette junction in transfected HEK293 cells through agarose gel electrophoresis. The PAX7-x2-fwd and pKER-detector-5’ primers were used in all reactions. The expected band size is 750 bp. Each lane represents one technical replicate for the given condition from the same experiment.

**Figure 5-figure supplement 1.**
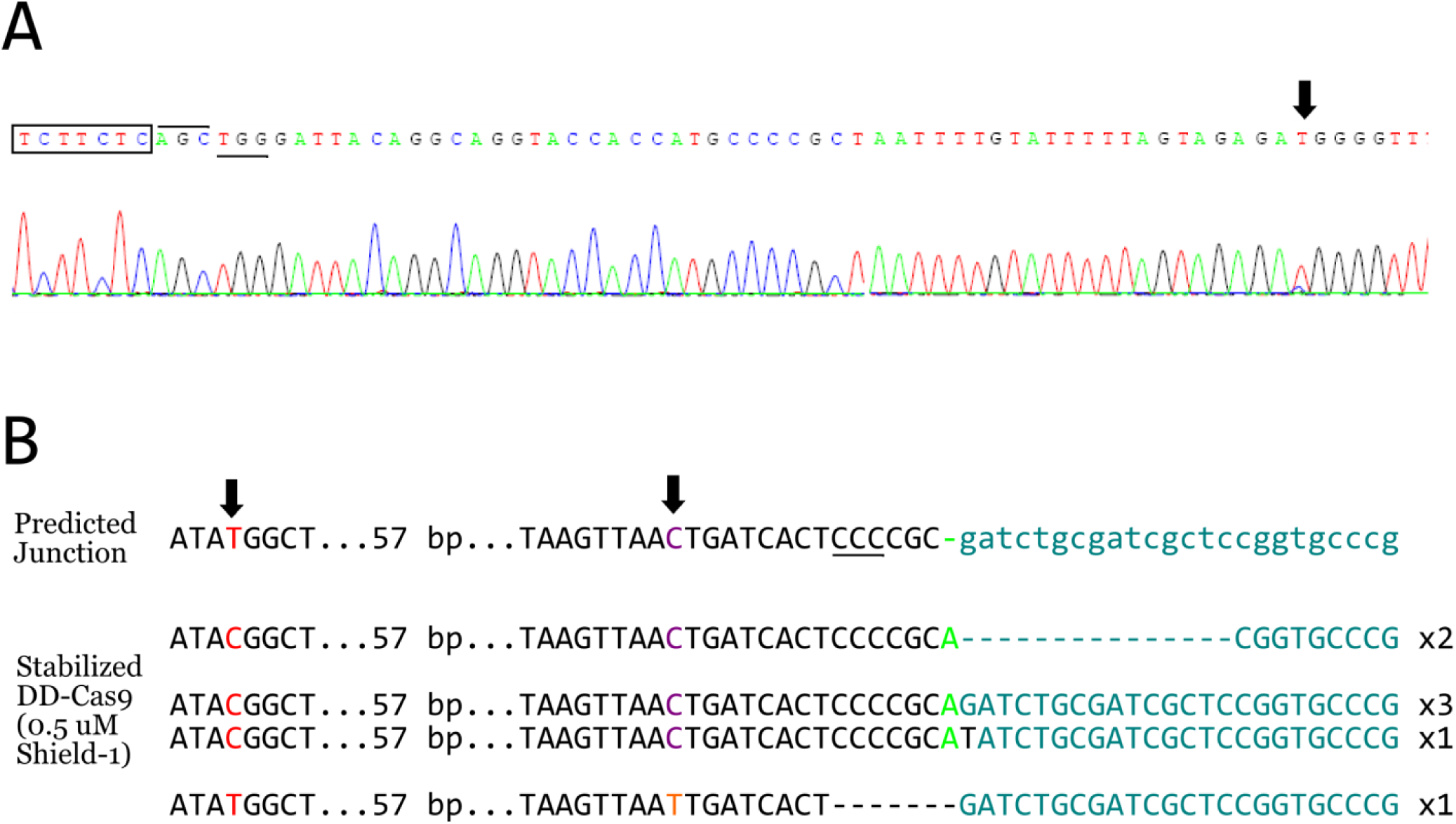
Further analysis of genome-cassette junctions resulting from KiBL at the H11 locus. (A) Chromatogram of bulk sequenced 3’pKER-Clover cassette-3’genome junctions resulting from the use of stabilized Cas9-DD. Boxed sequence is the 3’ end of the cassette. Overlined sequence is the expected first three bases of the 3’ sgRNA target site. Underlined sequence is the PAM. Arrowhead indicates the 3’ allele-specific SNP at this locus. (B) Sequenced amplicons of the 5’ genome-5’ pKER cassette junction from JF10 cells treated with the H11 PAMs Out sgRNA pair and stabilized DD-Cas9 (0.5 μM Shield-1). Left arrowhead indicates 5’ allele-specific SNP (in red). Right arrowhead indicates cytosine that differs from human reference sequence. PAM is underlined. Green denotes fourth base from the PAM. Turquoise denotes cassette sequence.

**Figure 5-figure supplement 2.**
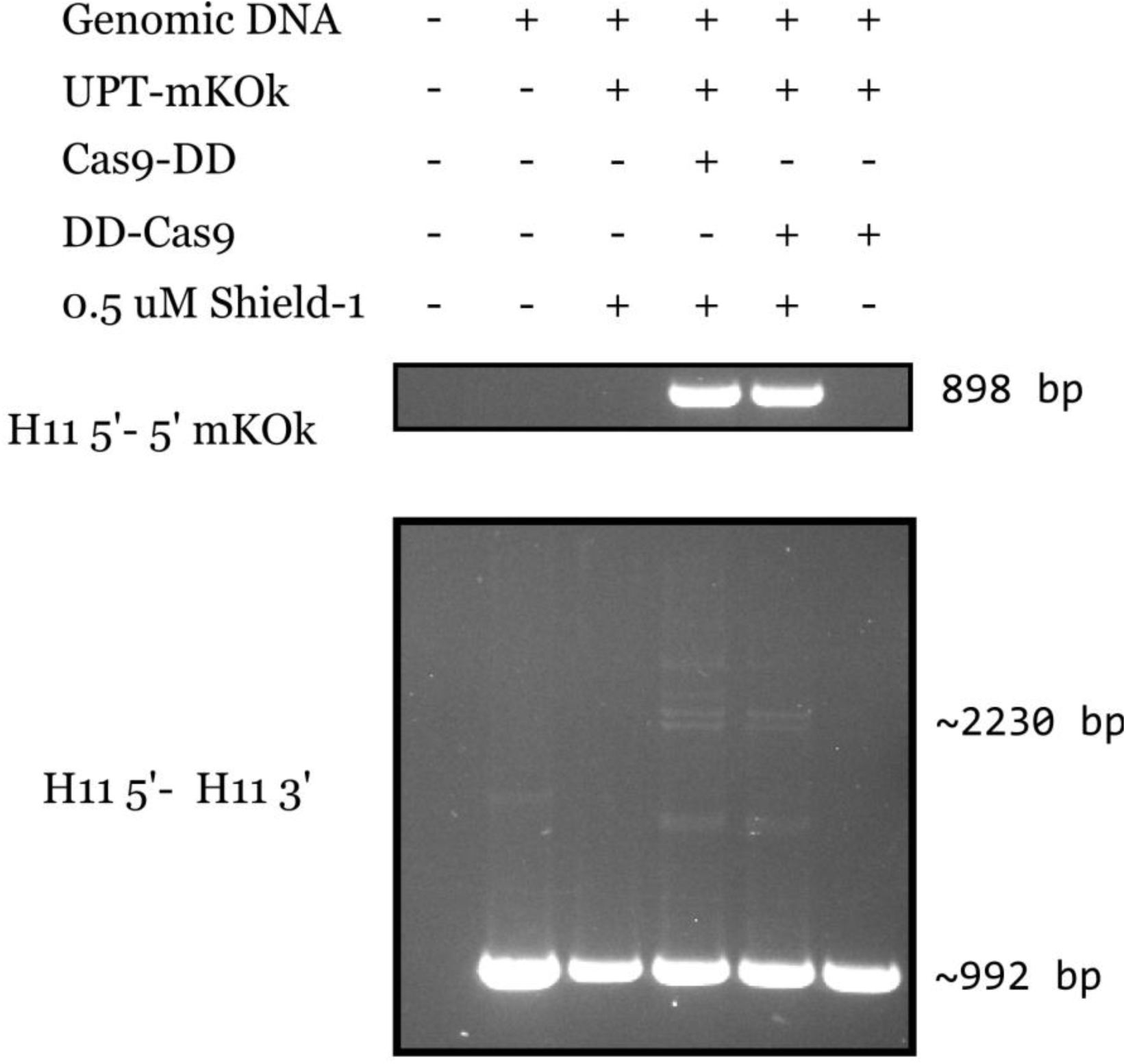
Comparison of the effect of destabilization of DD-Cas9 on KiBL. PCR was performed to analyze the H11 5’-5’ pKER cassette junction and the H11 locus of sorted JF10 hiPSCs transfected with pKER-mKOκ cassettes and the H11 PAMs Out sgRNA pair with Cas9-DD or DD-Cas9 in the presence or absence of Shield-1. The primers H11-x1-fwd and mKOκ-AS-N2 were used to amplify the H11 5’-5’ cassette junction and the primers H11-X1-fwd and H11-X1-rev were used to amplify the H11 locus. Gel electrophoresis was performed to visualize bands. The expected band size of the H11 5’-5’ mKOκ cassette junction is 898 bp. The expected band size of the unmodified H11 locus is 992 bp and the expected size of the KiBL allele is 2233 bp.

## References

Anders C, Niewoehner O, Duerst A, Jinek M. 2014. Structural basis of PAM-dependent target DNA recognition by the Cas9 endonuclease. Nature 513:569–73. doi:10.1038/nature13579.

Auer TO, Duroure K, De Cian A, Concordet, J-P, Del Bene F. 2014. Highly efficient CRISPR/Cas9-mediated knock-in in zebrafish by homology-independent DNA repair. Genome Research 24: 142–53. doi:10.1101/gr.161638.113.

Banaszynski LA, Chen, L-C, Maynard-Smith LA, Ooi AGL, Wandless TJ. 2006. A Rapid, reversible, and tunable method to regulate protein function in living cells using synthetic small molecules. Cell 126: 995–1004. doi:10.1016/j.cell.2006.07.025.

Banaszynski LA, Sellmeyer MA, Contag CH, Wandless TJ, Thorne SH. 2008. Chemical control of protein stability and function in living mice. Nature Medicine 14:1123–7. doi:10.1038/nm.1754.

Barrangou R, Fremaux C, Deveau H, Richards M, Boyaval P, Moineau S, Romero DA, Horvath P. 2007. CRISPR provides acquired resistance against viruses in prokaryotes. Science 315: 1709–12. doi:10.1126/science.1138140.

Barzel A, Paulk NK, Shi Y, Huang Y, Chu K, Zhang F, et al. 2015. Promoterless gene targeting without nucleases ameliorates haemophila B in mice. Nature 517: 360–4. doi:10.1038/nature13864.

Beerman I, Seita J, Inlay MA, Weissman IL, Rossi DJ. 2014. Quiescent hematopoietic stem cells accumulate DNA damage during aging that is repaired upon entry into the cell cycle. Cell Stem Cell 15: 37–50. doi:10.1016/j.stem.2014.04.016.

Bétermier M, Bertrand P, Lopez BS. 2014. Is non-homologous end-joining really an inherently error-prone process? PLOS Genetics 10:e1004086. doi:10.1371/journal.pgen.1004086.

Böttcher R, Hollman M, Merk K, Nitschko V, Obermaier C, Philippou-Massier J et al. 2014. Efficient chromosomal gene modification with CRISPR/*cas9* and PCR-based homologous recombination donors in cultured *Drosophila* cells. Nucleic Acids Research 42:e89. doi:10.1093/nar/gku289.

Brandsma I, van Gent DC. 2012. Pathway choice in DNA double strand break repair: observations of a balancing act. Genome Integrity 3: 9. doi:10.1186/2041-9414-3-9.

Byrne SM, Ortiz L, Mali P, Aach J, Church GM. 2014. Multikilobase homozygous targeted gene replacement in human induced pluripotent stem cells. Nucleic Acids Research. doi:10.1093/nar/gku1246.

Canver MC, Bauer DE, Dass A, Yien YY, Chung J, Masuda T et al. 2014. Characterization of genomic deletion efficiency mediated by CRISPR/Cas9 in mammalian cells. The Journal of Biological Chemistry. doi:10.1074/jbc.M114.564625.

Cencic R, Miura H, Malina A, Robert F, Ethier S, Schmeing TC, Dostie J, Pelletier J. 2014. Protospacer adjacent motif (PAM)-distal sequences engage CRISPR Cas9 DNA target cleavage. PLOS ONE 9:e109213. doi:10.1371/journal.pone.0109213.

Cho SW, Kim S, Kim Y, Kweon J, Kim HS, Bae S, Kim J-S. 2014. Analysis of off-target effects of CRISPR/Cas-derived RNA-guided endonucleases and nickases. Genome Research 24:132–41. doi:10.1101/gr.162339.113.

Chu J, Haynes RD, Corbel SY, Li P, González-González E, Burg JS, et al. 2014. Non-invasive intravital imaging of cellular differentiation with a bright red-excitable fluorescent protein. Nature Methods. doi:10.1038/nmeth.2888.

Chu VT, Weber T, Wefers B, Wurst W, Sander S, Rajewsky K, Kühn R. 2015. Increasing the efficiency of homology-directed repair for CRISPR-Cas9-induced precise gene editing in mammalian cells. Nature Biotechnology. doi:10.1038/nbt.3198.

Cong L, Ran FA, Cox D, Lin S, Barretto R, Habib N et al. 2013. Multiplex genome engineering using CRISPR/Cas9 sytems. Science 339:819–23. doi:10.1126/science.1231143.

Cristea S, Freyvert Y, Santiago Y, Holmes MC, Urnov FD Gregory PD, Cost GJ. 2013. In vivo cleavage of transgene donors promotes nuclease-mediated targeted integration. Biotechnology and Bioengineering 110:871–80. doi:10.1002/bit.24733.

Cuozzo C, Porcellini A, Angrisano T, Morano A, Lee B, Di Pardo A et al. 2007. DNA damage, homology-directed repair, and DNA methylation. PLoS Genetics 3: e110. doi:10.1371/journal.pgen.0030110.

Davis KM, Pattanayak V, Thompson DB, Zuris JA, Liu DR. 2015. Small molecule-triggered Cas9 protein with improved genome-editing specificity. Nature Chemical Biology AOP. doi:10.1038/nchembio.1793.

Dean FB, Hosono S, Fang L, Wu X, Faruqi AF, Bray-Ward P et al. 2002. Comprehensive human genome amplification using multiple displacement amplification. Proc. Nat. Acad. Sci. 99: 5261–6. doi:10.1073/pnas.082089499.

Demeter J, Lee SE, Haber JE, Stearns T. 2000. The DNA damage checkpoint signal in budding yeast is nuclear limited. Molecular Cell 6:487–92. doi:10.1016/S1097-2765(00)00047-2.

DiCarlo JE, Norville JE, Mali P, Rios X, Aach J, Church GM. 2013. Genome engineering in *Saccharomyces cerevisiae* using CRISPR-Cas systems. Nucleic Acids Research 41: 4336–43. doi:10.1093/nar/gkt135.

Farboud B, Meyer BJ. 2015. Dramatic enhancement of genome editing by CRISPR/Cas9 through improved guide RNA design. Genetics 199: 959–71. doi:10.1534/genetics.115.175166/-/DC1.

Findlay GM, Boyle EA, Hause RJ, Klein JC, Shendure J. 2014. Saturation editing of genomic regions by multiplex homology-directed repair. Nature 513:120–3. doi:10.1038/nature13695.

Flach J, Bakker ST, Mohrin M, Conroy PC, Pietras PC, Reynaud D et al. 2014. Replication stress is a potent driver of functional decline in ageing haematopoietic stem cells. Nature 512:198–202. doi:10.1038/nature13619.

Friedland AE, Tzur YB, Esvelt KM, Colaiácovo MP, Church GM, Calarco JA. 2013. Heritable genome editing in *C. elegans* via a CRISPR-Cas9 system. Nature Methods. doi:10.1038/nmeth.2532.

Frock RL, Hu J, Meyers RM, Hu, Y-J, Kii E, Alt FW. 2014. Genome-wide detection of DNA double-stranded breaks induced by engineered nucleases. Nature Biotechnology 33: 179–86. doi:10.1038/nbt.2916.

Fu BX, Hansen LL, Artiles KL, Nonet ML, Fire AZ. 2014. Landscape of target: guide homology effects on Cas9-mediated cleavage. Nucleic Acids Research 42:13778–87. doi:10.1093/nar/gku1102.

Fu Y, Foden JA, Khayter C, Maeder ML, Reyon D, Joung JK, Sander JD. 2013. High frequency off-target mutagenesis induced by CRISPR-Cas nucleases in human cells. Nature Biotechnology. doi:10.1038/nbt.2623.

Fu Y, Sander JD, Reyon D, Cascio VM, Joung JK. 2014. Improving CRISPR-Cas nuclease specificity using truncated guide RNAs. Nature Biotechnology 23: 279–84. doi:10.1038/nbt.2808.

Ghezraoui H, Piganeau M, Renouf B, Renaud, J-B, Sallmyr A, Ruis B. 2014. Chromosomal translocations in human cells are generated by canonical nonhomologous end-joining. Molecular Cell 55:1–14. doi:10.1016/j.molcel.2014.08.002.

Gratz SJ, Ukken FP, Rubinstein CD, Thiede G, Donohue LK, Cummings AM, O’Connor-Giles KM. 2014. Highly specific and efficient CRISPR/Cas9-catalyzed homology directed repair in *Drosophila*. Genetics 196: 961–71. doi:10.1534/genetics.113.160713/-/DC1.

Guilinger JP, Thompson DB, Liu DR. 2014. Fusion of catalytically inactive Cas9 to FokI nuclease improves the specificity of genome modification. Nature Biotechnology. doi:10.1038/nbt.2909

Harel I, Benayoun BA, Machado B, Singh PP, Hu, C-K, Pech MF et al. 2015. A platform for rapid exploration of aging and diseases in a naturally short-lived vertebrate. Cell 160: 1–14. doi:10.1016/j.cell.2015.01.038.

Heckl D, Kowalczyk MS, Yudovich D, Belizaire R, Puram RV, McConkey ME et al. 2014. Generation of mouse models of myeloid malignancy with combinatorial genetic lesions using CRISPR-Cas9 genome editing. Nature Biotechnology 22:941–6. doi:10.1038/nbt.2951.

Hsu PD, Scott DA, Weinstein JA, Ran FA, Konermann S, Agarwala V et al. 2013. DNA targeting specificity of RNA-guided Cas9 nucleases. Nature Biotechnology. doi:10.1038/nbt.2647.

Hu W, Kaminski R, Yang F, Zhang Y, Cosentino L, Li F et al. 2014. RNA-directed gene editing specifically eradicates and prevents new HIV-1 infection. Proc. Nat. Acad. Sci. doi:10.1073/pnas.1405186111.

Jinek M, Chylinski K, Fonfara I, Hauer M, Doudna JA, Charpentier E. 2012. A programmable dual-RNA-guided DNA endonuclease in adaptative bacterial immunity. Science 337:816–21. doi:10.1126/science.1225829.

Jinek M, East A, Cheng A, Lin S, Ma E, Doudna J. 2013. RNA-programmed genome editing in human cells. eLife 2: e00471. doi:10.7554/eLife.00471.

Kabadi AM, Ousterout DG, Hilton IB, Gersbach CA. 2014. Multiplex CRISPR/Cas9-based genome engineering from a single lentiviral vector. Nucleic Acids Research 42: e147. doi:10.1093/nar/gku749.

Kent WJ, Sugnet CW, Furrey TS, Roskin JM, Pringle TH, Zahler AM, Haussler D. 2002. The human genome browser at UCSC. Genome Research 12:996–1006. doi:10.1101/gr229102.

Kim D, Bae S, Park J, Kim E, Kim S, Yu HR et al. 2015. Digenome-seq: genome-wide profiling of CRISPR-Cas9 off-target effects in human cells. Nature Methods 12:237–42. doi:10.1038/nmeth.3248.

Kim H, Ishidate T, Ghanta KS, Seth M, Conte Jr. D, Shirayama M, Mello CC. 2014. A Co-CRISPR strategy for efficient genome editing in *Caenorhabditis elegans*. Genetics 197:1069–80. doi:10.1534/genetics.114.166389.

Kuscu C, Arslan S, Singh R, Thorpe J, Adli M. 2014. Genome-wide analysis reveals characteristics of off-target sites bound by the Cas9 endonuclease. Nature Biotechnology 32:677–83. doi:10.1038/nbt.2916

Lam AJ, St-Pierre F, Gong Y, Marshall JD, Cranfill PJ, Baird MA et al. 2012. Improving FRET dynamic range with bright green and red fluorescent proteins. Nature Methods 9:1005–12. doi:10.1038/nmeth.2171.

Li K, Wang G, Andersen T, Zhou P, Pu WT. 2014. Optimization of genome engineering approaches with the CRISPR/Cas9 system. PLOS ONE 9: e105779. doi:10.1371/journal.pone.0105779.

Lin S, Staahl BT, Alla RK, Doudna JA. 2014. Enhanced homology-directed human genome engineering by controlling timing of CRISPR/Cas9 delivery. eLife 3:e04766. doi: 10.7554/eLife.04766.

Lin Y, Cradick TJ, Brown MT, Deshmukh H, Ranjan P, Sarode N et al. 2014. CRISPR/Cas9 systems have off-target activity with insertions or deletions between target DNA and guide RNA sequences. Nucleic Acids Research 42:7473–85. doi:10.1093/nar/gku402.

Mali P, Yang L, Esvelt KM, Aach J, Guell M, DiCarlo JE et al. 2013a. RNA-guided human genome engineering via Cas9. Science 339: 823–6. doi:10.1126/science.1232033.

Mali P, Aach J, Stranges PB, Esvelt KM, Moosburner M, Kosuri S, Yang L, Church GM. 2013b. CAS9 transcriptional activators for target specificity screening and paired nickases for cooperative genome engineering. Nature Biotechnology. doi:10.1038/nbt.2675.

Mandal PK, Ferreira LMR, Collins R, Meissner TB, Boutwell CL, Friesen M et al. 2014. Efficient ablation of genes in human hematopoietic stem and effector cells using CRISPR/Cas9. Cell Stem Cell 15:643–52. doi:10.1016/j/stem/2014.10.004.

Maresca M, Lin VG, Guo N, Yang Y. 2013. Obligate ligation-gated recombination (ObLiGaRe): custom-designed nuclease-mediated targeted integration through nonhomologous end joining. Genome Research 23: 539–46. doi:10.1101/gr.145441.112.

Maruyama T, Dougan SK, Truttman MC, Bilate AM, Ingram JR, Ploegh HL. 2015. Nature Biotechnology. doi:10.1038/nbt.3190.

Maynard-Smith LA, Chen LC, Banaszynski LA, Ooi AG, Wandless TJ. 2007. A directed approach for engineering conditional protein stability using biologically silent small molecules. The Journal of Biological Chemistry 282:24866–72. doi:10.1074/jbc.M703902200.

Miyaoka Y, Chan AH, Judge LM, Yoo J, Huang M, Nguyen TD, Lizarraga PP, So, P-L, Conklin BR. 2014. Isolation of single-base genome-edited human iPS cells without antibiotic selection. Nature Methods 11: 291–3. doi:10.1038/nmeth.2840.

Nakade S, Tsubota T, Sakane Y, Kume S, Sakamoto N, Obara M, Daimon T et al. 2014 Microhomology-mediated end-joining-dependent integration of donor DNA in cells and animals using TALENs and CRISPR/Cas9. Nature Communications 5: 5560. doi:10.1038/ncomms6560.

Ousterout DG, Kabadi AM, Thakore PI, Majoros WH, Reddy TE, Gersbach CA. 2015. Multiplex CRISPR/Cas9-based genome editing for correction of dystrophin mutations that cause Duchenne muscular dystrophy. Nature Communications 6:6244. doi:10.1038/ncomms7244.

Pattanayak V, Lin S, Guilinger JP, Ma E, Doudna JA, Liu DR. 2013. High-throughput profiling of off-target DNA cleavage reveals RNA-programmed Cas9 nuclease specificity. Nature Biotechnology 31: 839–43. doi:10.1038/nbt.2673.

Platt RJ, Chen S, Zhou Y, Yim MJ, Swiech L, Kempton HR et al. 2014. CRISPR-Cas9 knockin mice for genome editing and cancer modeling. Cell 159:440–55. doi:10.1016/j.cell.2014.09.014.

Pruett-Miller SM, Reading DW, Porter SN, Porteus MH. 2009. Attenuation of zinc finger nuclease toxicity by small-molecule regulation of protein levels. PLoS Genetics 5: e1000376. doi:10.1371/journal.pgen.1000376.

Ramsahoye BH, Biniszkiewicz D, Lyko F, Clark V, Bird AP, Jaenisch R. 2000. Non-CpG methylation is prevalent in embryonic stem cells and may be mediated by DNA methyltransferase 3a. Proc. Nat. Acad. Sci. 97: 5237–42. doi:10.1073/pnas.97.10.5237.

Ran FA, Hsu PD, Lin, C-Y, Gootenberg JS, Konermann S, Trevino AE et al. 2013. Double-nicking by RNA-guided CRISPR Cas9 for enhanced genome editing specificity. Cell 154: 1380–9. doi:10.1016/j.cell.2013.08.021.

Rizzo MA, Springer GH, Granada B, Piston DW. 2004. An improved cyan fluorescent protein variant useful for FRET. Nature Biotechnology 22:445–9. doi:10.1038/nbt945.

Sadelain M, Papapetrou EP, Bushman FD. 2011. Safe harbours for the integration of new DNA in human genome. Nature Reviews Cancer. doi:10.1038/nrc3179.

Sternberg SH, Redding S, Jinek M, Greene EC, Doudna JA. 2014. DNA interrogation by the CRISPR RNA-guided endonuclease Cas9. Nature 507: 62–7. doi:10.1038/nature13011.

Swiech L, Heidenreich M, Banerjee A, Habib N, Li Y, Trombetta J, Sur M, Zhang F. 2015 *In vivo* interrogation of gene function in the mammalian brain using CRISPR-Cas9. Nature Biotechnology 33: 102–106. doi:10.1038/nbt.3055.

Tasic B, Hippenmeyer S, Wang C, Gamboa M, Zong H, Chen-Tsai Y, Luo L. 2011. Site-specific integrase-mediated transgenesis in mice via pronuclear injection. Proc. Nat. Acad. Sci. 108: 7902–7. doi:10.1073/pnas.1019507108.

Tsai SQ, Wyvekens N, Khayter C, Foden JA, Thapar V, Reyon D et al. 2014a. Dimeric CRISPR RNA-guided FokI nucleases for highly specific genome editing. Nature Biotechnology. doi:10.1038/nbt.2908.

Tsai SQ, Zheng Z, Nguyen NT, Liebers M, Topkar VV, Thapar V et al. 2014b. GUIDE-seq enables genome-wide profiling of off-target cleavage by CRISPR-Cas nucleases. Nature Biotechnology. doi:10.1038/nbt.3117.

Tsutsui H, Karasawa S, Okamura Y, Miyawaki A. 2008. Improving membrane voltage measurements using FRET with new fluorescent proteins. Nature Methods 5:683–5. doi:10.1038/nmeth.1235.

Vahidi Ferdousi L, Rocheteau P, Chaynot R, Montagne B, Chaker Z, Flamant P, Tajbakhsh S, Ricchetti M. 2014. More efficient repair of DNA double-strand breaks in skeletal muscle stem cells compared to their committed progeny. Stem Cell Research 13: 492–507. doi:10.1016/j.scr.2014.08.005.

van Heemst D, Brugmans L, Verkaik NS, van Gent DC. 2004. End-joining of blunt DNA double-strand breaks in mammalian fibroblasts is precise and requires DNA-PK and XRCC4. DNA Repair 3: 43–50. doi:10.1016/j.dnarep.2003.09.004.

Veres A, Gosis BS, Ding Q, Collins R, Ragavendran A, Brand H et al. 2014. Low incidence of off-target mutations in individual CRISPR-Cas9 and TALEN targeted human stem cell clones detected by whole-genome sequencing. Cell Stem Cell 15: 27–30. doi:10.106/j.stem.2014.04.020.

Wang J, Quake SR. 2014. RNA-guided endonuclease provides a therapeutic strategy to cure latent herpesviridae infection. Proc. Nat. Acad. Sci. 111:13157–62. doi:10.1073/pnas1410785111.

Wang X, Wang W, Wu X, Wang J, Wang Y, Qiu Z et al. 2015. Unbiased detection of off-target cleavage by CRISPR-Cas9 and TALENs using integrase-defective lentiviral vectors. Nature Biotechnology 33:175–8. doi:10.1038/nbt.3127.

Wright AV, Sternberg SH, Taylor DW, Staahl BT, Bardales JA, Kornfeld JE, Doudna JA. 2015. Rational design of a split-Cas9 enzyme complex. Proc. Nat. Acad. Sci. 110:2984–9. doi:10.1073/pnas.1501698112.

Wu X, Scott DA, Kriz AJ, Chiu AC, Hsu PD, Dadon DB et al. 2014. Genome-wide binding of the CRISPR endonuclease Cas9 in mammalian cells. Nature Biotechnology. doi:10.1038/nbt.2889.

Xie F, Ye L, Chang JC, Beyer AI, Wang J, Muench MO, Kan YW. 2014. Seamless gene correction of β-thalassemia mutations in patient-specific iPSCs using CRISPR/Cas9 and *piggyBac*. Genome Research. doi:10.1101/gr.173427.114.

Xue W, Chen S, Yin H, Tammela T, Papagiannakopoulos T, Joshi NS, Cai W, Yang G et al. 2014. CRISPR-mediated direct mutation of cancer genes in the mouse liver. Nature 514: 380–4. doi:10.1038/nature13589.

Yang H, Wang H, Shivalila CS, Cheng AW, Shi L, Jaenisch R. 2013. One-step generation of mice carrying reporter and conditional alleles by CRISPR/Cas-mediated genome engineering. Cell 154: 1370–9. doi:10.1016/j.cell.2013.08.022.

Yang L, Guell M, Bryne S, Yang JL, De Los Angeles A, Mali P, et al. 2013. Optimization of scarless human stem cell genome editing. Nucleic Acids Research. doi:10.1093/nar/gkt555.

Yang L, Grishin D, Wang G, Aach J, Zhang, C-Z, Chari R et al. 2014. Targeted and genome-wide sequencing reveal single nucleotide variations impacting specificity of Cas9 in human stem cells. Nature Communications 5: 5507. doi:10.1038/ncomms6507.

Zhang X, Koolhaas WH, Schnorrer F. 2014. A versatile two-step CRISPR-and RMCE-based strategy for efficient genome engineering in *Drosophila*. G3: Genes | Genomes | Genetics 4:2409–18. doi:10.1534/g3.114.013979/-/DC1.

Zhang Y, Vanoli F, LaRocque JR, Krawczyk PM, Jasin M. 2014. Biallelic targeting of expressed genes in mouse embryonic stem cells using the Cas9 system. Methods 69:171–8. doi:10.1016/j.ymeth.2014.05.003.

Zheng Q, Cai X, Tan MH, Schaffert S, Arnold CP, Gong X et al. 2014. Precise gene deletion and replacement using the CRISPR/Cas9 system in human cells. Biotechniques 57: 115–24. doi:10.2144/000114196.

Zhou J, Wang J, Shen B, Chen L, Su Y, Yang J et al. 2014. Dual sgRNAs facilitate CRISPR/Cas9-mediated mouse genome targeting. FEBS Journal. doi:10.1111/febs.12735.

Zhu F, Gamboa M, Farruggio AP, Hippenmeyer S, Tasic B, Schüle B et al. 2014. DICE, an efficient system for iterative genomic editing in human pluripotent stem cells. Nucleic Acids Research. doi:10.1093/nar/gkt/1290.

Zhu Z, Gonzalez F, Huangfu D. 2014. The iCRISPR platform for rapid genome editing in human pluripotent stem cells. Methods in Enzymology 546:215–50. doi:10.1016/B978-0-12-801185-0.00011-8.

